# Transcriptional and Cellular Diversity of the Human Heart

**DOI:** 10.1101/2020.01.06.896076

**Authors:** Nathan R. Tucker, Mark Chaffin, Stephen J. Fleming, Amelia W. Hall, Victoria A. Parsons, Kenneth Bedi, Amer-Denis Akkad, Caroline N. Herndon, Alessandro Arduini, Irinna Papangeli, Carolina Roselli, François Aguet, Seung Hoan Choi, Kristin G. Ardlie, Mehrtash Babadi, Kenneth B. Margulies, Christian M. Stegmann, Patrick T. Ellinor

## Abstract

**Introduction:** The human heart requires a complex ensemble of specialized cell types to perform its essential function. A greater knowledge of the intricate cellular milieu of the heart is critical to increase our understanding of cardiac homeostasis and pathology. As recent advances in low input RNA-sequencing have allowed definitions of cellular transcriptomes at single cell resolution at scale, here we have applied these approaches to assess the cellular and transcriptional diversity of the non-failing human heart.

**Methods:** Microfluidic encapsulation and barcoding was used to perform single nuclear RNA sequencing with samples from seven human donors, selected for their absence of overt cardiac disease. Individual nuclear transcriptomes were then clustered based upon transcriptional profiles of highly variable genes. These clusters were used as the basis for between-chamber and between-sex differential gene expression analyses and intersection with genetic and pharmacologic data

**Results:** We sequenced the transcriptomes of 287,269 single cardiac nuclei, revealing a total of 9 major cell types and 20 subclusters of cell types within the human heart. Cellular subclasses include two distinct groups of resident macrophages, four endothelial subtypes, and two fibroblasts subsets. Comparisons of cellular transcriptomes by cardiac chamber or sex reveal diversity not only in cardiomyocyte transcriptional programs, but also in subtypes involved in extracellular matrix remodeling and vascularization. Using genetic association data, we identified strong enrichment for the role of cell subtypes in cardiac traits and diseases. Finally, intersection of our dataset with genes on cardiac clinical testing panels and the druggable genome reveals striking patterns of cellular specificity.

**Conclusions:** Using large-scale single nuclei RNA sequencing, we have defined the transcriptional and cellular diversity in the normal human heart. Our identification of discrete cell subtypes and differentially expressed genes within the heart will ultimately facilitate the development of new therapeutics for cardiovascular diseases.

## Introduction

The heart is an organ that acts without rest, ceaselessly beating over 2 billion times in the average human lifetime. Given the heart’s central function as a pump, it is understandable that much of the cardiac research focus has been centered on the cell subtype most responsible for contractile functionality, the cardiomyocyte. However, cardiomyocytes do not function in isolation, instead contracting as part of a complex ensemble of specialized cell types including those responsible for tissue perfusion, remodeling of the interstitial space, and autonomic regulation. A greater understanding of the complex cellular milieu of the heart is critical to advance our understanding of cardiac homeostasis and pathology.

Analysis of transcription of RNA species, a highly dynamic process, is one method for defining cell types and states. To date, transcriptional analyses of the human heart have largely been performed in bulk tissue RNA sequencing studies. While these studies have yielded important insight into regional and pathological differences in tissue-level expression, they are unable to resolve the cell types from which any differential expression occurs. Recent advances in single cell RNA sequencing, particularly technologies centered on microfluidic encapsulation and cellular barcoding [1, 2] have made deconvolution of these expression profiles technologically feasible. Large efforts are currently underway to define the cellular diversity in all organ systems. Among these, the Human Cell Atlas (HCA) [3] and Human BioMolecular Atlas Program (HuBMAP, https://commonfund.nih.gov/hubmap) are of particular note in humans, while the *Tabula Muris* project [4] has provided valuable insight into the murine cell subtype transcriptome. Due to challenges with tissue availability and cellular isolation, there have been relatively few studies of the cardiac system to date. Some recent analyses of heart tissue from humans [5, 6] and model systems [4, 7] have recently been published, but are limited in scope. Thus, a comprehensive analysis of cell subtype expression profiles from the non-failing human heart has yet to be performed. The transcriptional map of the non-failing human heart at single-cell resolution, together with an understanding of its normal inter-individual variability, crucially serves as a baseline against which one can obtain equally high-resolution and quantitative maps of cardiac pathologies.

In the presently described study, we perform single nuclear RNA-sequencing (snRNAseq) on 287,269 nuclei derived from the four chambers of the normal human heart. We identified 9 major cell types and more than 20 cell subtypes. We observed marked differences in cell subtype transcription by chamber, laterality, and gender. We then intersected the snRNAseq data with the results from genome wide association studies to prioritize cell subtypes for cardiovascular disease risk and with the druggable genome to facilitate the identification of novel therapeutic targets for cardiovascular diseases. Finally, our data provides a methodological framework and large-scale resource available to the broader scientific community.

## Results

### Single-nucleus RNA-sequencing of the human adult myocardium

We obtained cardiac tissue samples from seven potential transplant donors, including four women and three men, without any clinical evidence of cardiac dysfunction (Table 1). Tissue samples taken from the lateral aspect of the four cardiac chambers were subjected to nuclear isolation and processing for single nucleus RNA-sequencing (10x Genomics 3’ Single Cell Solution v2). Each sample was processed in replicate, and the second sample underwent a modification in reverse transcription that significantly increased library complexity (**Methods**). In total, 56 libraries were generated which were then subjected to cell calling, background adjustment, quality control filtering and cell alignment (**Methods**). The workflow for filtration steps and resultant values of samples or cells passing QC at various phases are contained within **Figure S1A**.

**Table 1:** Clinical characteristics of transplant donors

| File # | Sex | Age (yr) | Weight (kg) | Height (cm) | Heart Weight (g) | LV Mass (g) | LVMI(g/m<sup>2</sup>) | LVEDD (cm) | LVESD (cm) | PW Thick (cm) | LVEF (%) | Creat (mg/dl) |
| --- | --- | --- | --- | --- | --- | --- | --- | --- | --- | --- | --- | --- |
| P1221 | Female | 52 | 59 | 156 | 300 |  |  | 4.2 | 2.9 | 0.8 | 75 | 0.7 |
| P1600 | Female | 51 | 68 | 162.6 | 213 | 134 | 76.5 | 4.2 | 2.8 | 0.7 | 50 | 0.8 |
| P1666 | Male | 54 | 62 | 173 | 262 | 159 | 92.1 |  |  | 1.2 | 65 | 0.75 |
| P1681 | Male | 39 | 61 | 170 | 400 | 232 | 136.7 | 4.5 | 3 | 0.9 | 60 | 0.7 |
| P1702 | Male | 59 | 62 | 177 | 386 | 206 | 118.0 | 4.6 | 2.7 | 0.8 | 60 | 1.38 |
| P1708 | Female | 60 | 63 | 160 | 281 | 159 | 95.0 | 3.9 | 1.8 | 1.1 | 65 | 0.47 |
| P1723 | Female | 47 | 79 | 167 | 310 | 205 | 107.1 |  |  | 0.9 | 60 | 0.8 |

LV: Left ventricle, LVMI: left ventricular mass index, LVEDD: left ventricular end diastolic dimension, LVESD: left ventricular end systolic dimension, PW Thick: posterior wall thickness, LVEF: left ventricular ejection fraction, Creat: creatinine

In total, 287,269 cells from 44 libraries were utilized in downstream analyses, including identification of cell types and states (**Figure S1B**, **Table ST1**). When constructing transcriptional maps of human donors, we used single-cell variational inference (*scVI*) batch correction to prevent cells from segregating by individual donors within cell type clusters (**Figure S2A**). Additionally, use of CellBender *remove-background* allowed for calling of cells with lower transcriptional complexity, especially in the context of the relatively complex cardiomyocyte nuclei (**Figure S2B**), while also removing the contamination from ambient mRNAs. Importantly, a 3’ capture-derived RNA sequencing library is designed to capture poly-adenylated transcripts and thus does not completely identify the RNA molecules present within a ribosomal RNA-depleted, fragment-based, bulk RNA sequencing experiment.

A total of 17 distinct cell clusters were observed following unsupervised Louvain clustering at a resolution of 1.0. Distributions of cell clusters by chamber specific UMAP representations are shown in Figure 1A which are combined within a global UMAP representation in Figure 1B. We were able to group these into 9 major cell types by canonical marker and ontology analysis, followed by analyses of cell type substructure within each of these groups. Cell clusters are well represented across samples with a few notable exceptions (Figure 1C). First, cardiomyocytes derived from the atria cluster independently of those from the ventricle. Second, one ventricular cardiomyocyte cluster is largely found in the right ventricle of a single sample, **P1708**. Third, lymphocytes were preferentially found in the left ventricle of sample **P1723**. In addition, we believe two specific clusters represent cytoplasmic fragments as they are enriched for reads mapping to mature transcripts and mitochondrial genes (**Figure S2C,D**). The following sections will detail the features of each cell cluster, which are described by marker genes in Figure 2 and **Table ST2** and analyzed for gene ontology biological function terms in Figure 2. Markers genes were determined as those which display an area under the receiver operating characteristic curve (AUC) value of greater than 0.7 and an average natural log fold change greater than 0.6 (**Methods**). In cases when an insufficient number of genes was identified to define a cluster, additional genes with lower levels of overall expression, but strong selectivity for the target cluster of interest, were used for cell type definitions. These were defined as genes expressed in at least 5% of target cells and with a standardized positive predictive value (PPV50) greater than 0.90 (**Methods**). For subclustering analyses, a similar approach was employed but lowering the threshold for marker genes to an AUC greater than 0.65 and average natural log fold change greater than 0.5. As with the clusters from the global map, for some subclusters additional genes expressed in at least 5% of cells in the target subcluster with PPV50 greater than 0.90 were interrogated to assign subcluster labels (**Methods**).

**Figure 1:**
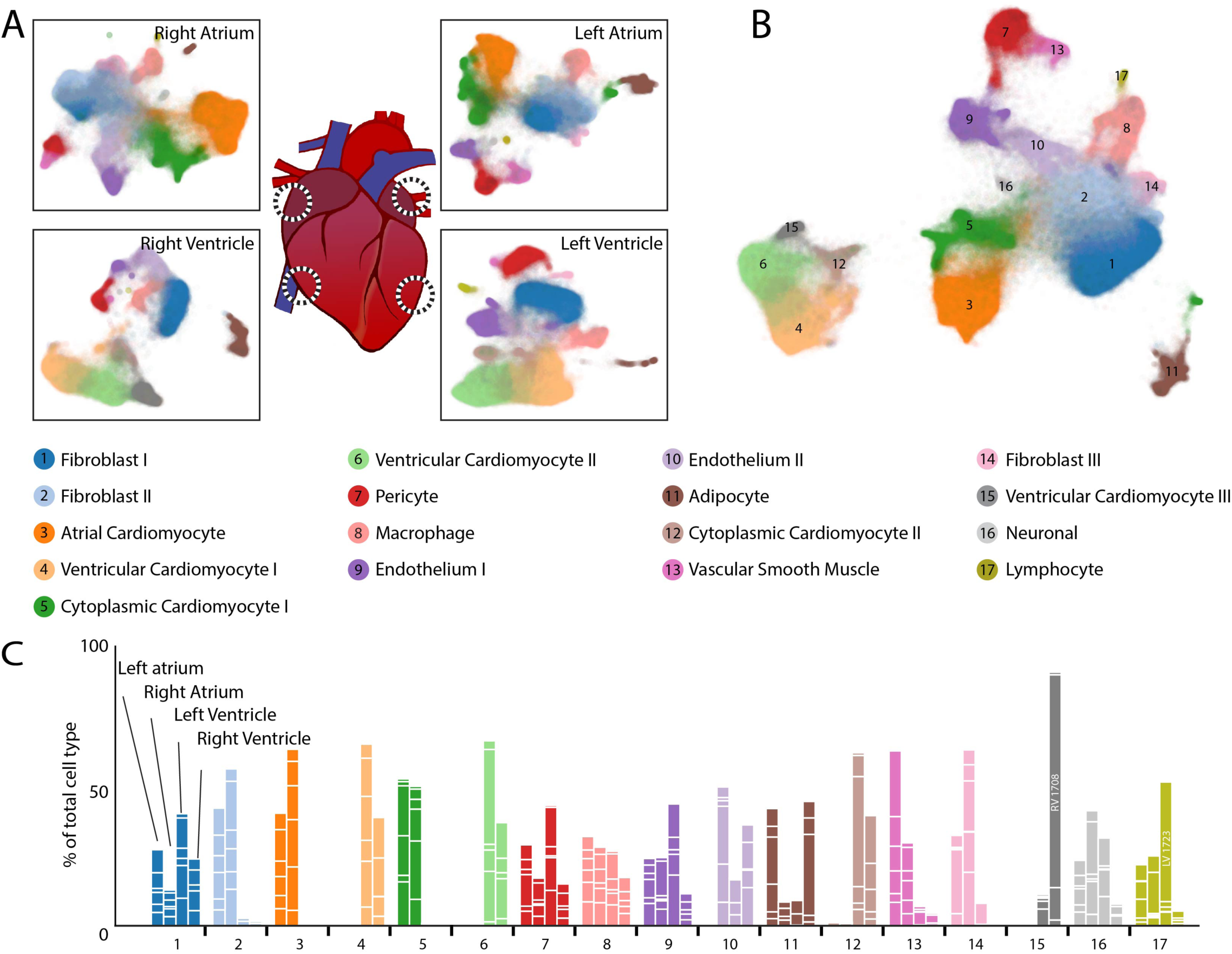
Observed cell types in the adult human heart. A: UMAP plot displaying cellular diversity present in the human heart by chamber. Each dot represents an individual cell. Colors correspond to the cell cluster labels below the panel. B: Combined UMAP plot containing a total of 287,269 cells from 7 individuals. Colors and numbers correspond to the cell cluster labels as listed in the lower panel. C: Relative representation of cell clusters by sample. Aggregation of four bars for each cell cluster equals 100% for each cell type. White lines within bars separate individual sample contributions. Colors correspond to the cell type descriptions displayed in the panel above.

**Figure 2:**
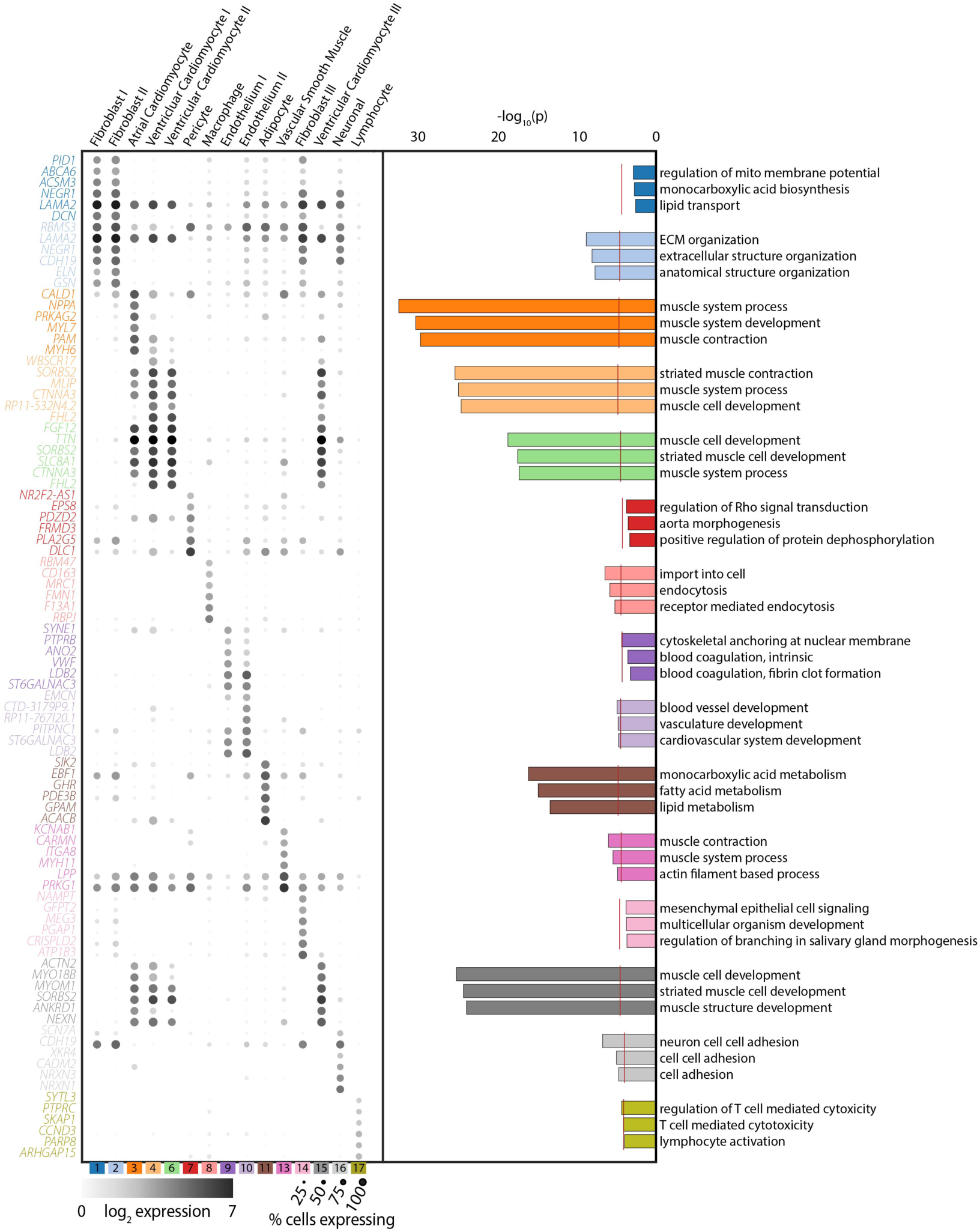
Gene and ontology definitions of observed cardiac cell clusters. Left panel: Dot plots display the top 6 marker genes for each supercluster as determined by AUC. The size of the dot represents the percentage of cells within the cluster where each marker is detected while the gradation corresponds to the mean log_2_ of the counts normalized by total counts per cell times 10,000. Right panel: Gene ontology enrichment analysis as performed by GOStats using all genes which reach an AUC threshold of greater than 0.70 and an average log fold-change greater than 0.60 for the given cell cluster. Red dotted line indicates a Bonferroni statistical significance threshold. The top three gene ontologies are shown for each cell cluster.

### Nine major cell types and more than twenty subclusters of cell types in the human heart

#### Distinct transcriptional profile of atrial and ventricular cardiomyocytes

Cell clusters 3, 4, 5, 6, 12, and 15 comprise the most frequent major cell type of cardiomyocytes and reflect strong expression of genes involved in canonical excitation-contraction function. Clusters 5 and 12 displayed an enrichment of mature mRNAs (**Figure S2D**), suggesting that non-nuclear regions were the source of these “cells.” We removed these clusters from subsequent analyses as the clear differences between cytoplasm and nuclei would further confound comparisons across chamber and sex. After this exclusion, cardiomyocytes represented 35.9% of observed cells. Cluster 3 displayed canonical markers of the atrium, including *NPPA* (AUC_3_=0.91), *MYL7* (AUC_3_=0.93) and *MYH6* (AUC_3_=0.96) (Figure 2 and **Table ST2**). Clusters 4, 6, and 15 displayed obvious markers of mature cardiomyocytes such as *TTN* (AUC_4_=0.85, AUC_6_=0.86, AUC_15_=0.79) and *MYH7* (AUC_4_=0.87, AUC_15_=0.79), but had fewer known markings of ventricular specificity in the global analysis (Figure 2 and **Table ST2**). This is likely due to the splitting of ventricular cardiomyocytes amongst multiple clusters by the Louvain algorithm such that some subclusters are included in the reference group for marker gene identification in a given cluster. A separate analysis of atrial versus ventricular cardiomyocytes resolved this issue and is discussed in the cross chamber comparisons below.

Subclustering of aggregated cardiomyocytes reveals 5 subclusters (**Figure S3A**). Cardiomyocyte subcluster 1 (CM-S1) corresponds to cluster 3 from the global map and contains all atrial cardiomyocytes. Within the ventricular cardiomyocytes, cluster CM-S5 has enrichment for mitochondrial components and an increased mature transcript proportion suggesting these may also be cytoplasmic contaminants. CM-S4 correlates strongly to cluster 15 in the global map and displays increased expression of *ANKRD1* (AUC_CM-S4_=0.82), which is thought to have a role in cardiomyopathy associated remodeling [8] and *KCP* (PPV50_CM-S4_=0.91), a BMP modifier whose expression is associated with heart failure [9] (**Figure S3A** and **Table ST3**). These cells are most often found in the right ventricle of a single donor (73% from **P1708**), and may represent a marker of a sub-clinical cardiac pathology.

#### Identification of activated and non-activated cardiac fibroblasts

By volume, cardiomyocytes comprise the majority of heart mass; however, in the absence of structural heart disease, fibroblast are roughly equivalent to cardiomyocytes in cell number. As the hearts used in this study were largely free of fibrotic remodeling (**Figure S4A**), we expected similar representation for fibroblasts and cardiomyocyte nuclei within our data. The cells from the combination of clusters 1, 2, and 14 represent cardiac fibroblasts, constituting 32.4% of observed cells. These cells display common markers of fibroblast lineages, with enriched expression of known fibroblast genes such as *DCN* (AUC_1_=0.85, AUC_2_=0.83), which encodes the proteoglycan decorin which regulates collagen fibrillogenesis, and (*ELN* (AUC_1_=0.71, AUC_2_=0.86), which produces elastin, a major component of the extracellular matrix (Figure 2 and **Table ST2**). The former was used to evaluate the distribution of the fibroblasts in our tissue samples, which exhibit the traditional interstitial localization observed in previous work (**Figure S4B**). In addition to extracellular matrix proteins, members of the ATP binding cassette subfamily of transmembrane transporters, including *ABCA6*, *-8* and *-9*, were also preferentially expressed in one or more of these clusters (*ABCA6*: AUC_1_=0.80, AUC_2_=0.77; *ABCA8*: AUC_1_=0.79, AUC_2_=0.79, *ABCA9*: AUC_1_=0.75, AUC_2_=0.74) **(Table ST2)**. Analysis of ontology for specific genes in this class display expected terms in the realm of extracellular matrix and structural organization, with the greatest enrichment in cluster 2 (Figure 2). No terms reached significance thresholds for clusters 1 and 14, perhaps as a consequence of a lower number of genes surpassing our criteria of a marker gene within these clusters (15 and 48, respectively). This is largely a consequence of including other fibroblast clusters in the reference outgroup for marker gene testing.

To further evaluate the structure within the fibroblast population, we performed local clustering of these cells, from which 4 populations were observed (Figure 3A). Importantly, subcluster FB-S2, which composes a large proportion of cluster 2 in the global map, shows an enrichment for *NPPA*, a known marker of atrial cardiomyocytes (Figure 3B). Whether this is truly fibroblast specific *NPPA* expression, an artifact derived from cardiomyocyte/fibroblast nuclear doublets, or a result of the presence of *NPPA* transcript in the extranuclear contaminant, requires further investigation. Cluster FB-S3 displays enriched expression of fibrosis associated genes *NOX4* (AUC_FB-S3_=0.70) and *IGF1* (AUC_FB-S3_=0.69), and cluster FB-S4, which corresponds to cluster 14 in the main map, exhibits clear upregulation of pro-fibrotic markers, including *ADAMTS4* (AUC_FB-S4_=0.69), which encodes a pro-fibrotic metalloprotease, *VCAN* (AUC_FB-S4_=0.69), which encodes the proteoglycan versican [10], and *AXL* (AUC_FB-S4_=0.69), which encodes a receptor tyrosine kinase associated with pathologic remodeling [11](Figure 3B**, Table ST3**). Further interrogation of these cells via RNA *in situ* hybridization with α-*ADAMTS4-*specific probes demonstrates an interstitial distribution throughout the tissue rather than being localized to a particular region (Figure 3C), suggesting that an organ wide event stimulated this fibroblast state transition. To attempt to identify the lineage of these fibroblast subclusters, we intersected our data with those from fibroblast activation in mice and humans.[12, 13] None of these clusters are enriched for expression of canonical markers for fibroblast activation (*POSTN*), myofibroblast transition (*MYH11*, *FAP*), or transformation to fibrocytes (*CHAD*, *COMP*) (Figure 3B). Whether these cells are a previously undefined state in canonical fibroblast activation, or are instead an entirely non-canonical form of fibroblast will be the focus of future work.

**Figure 3:**
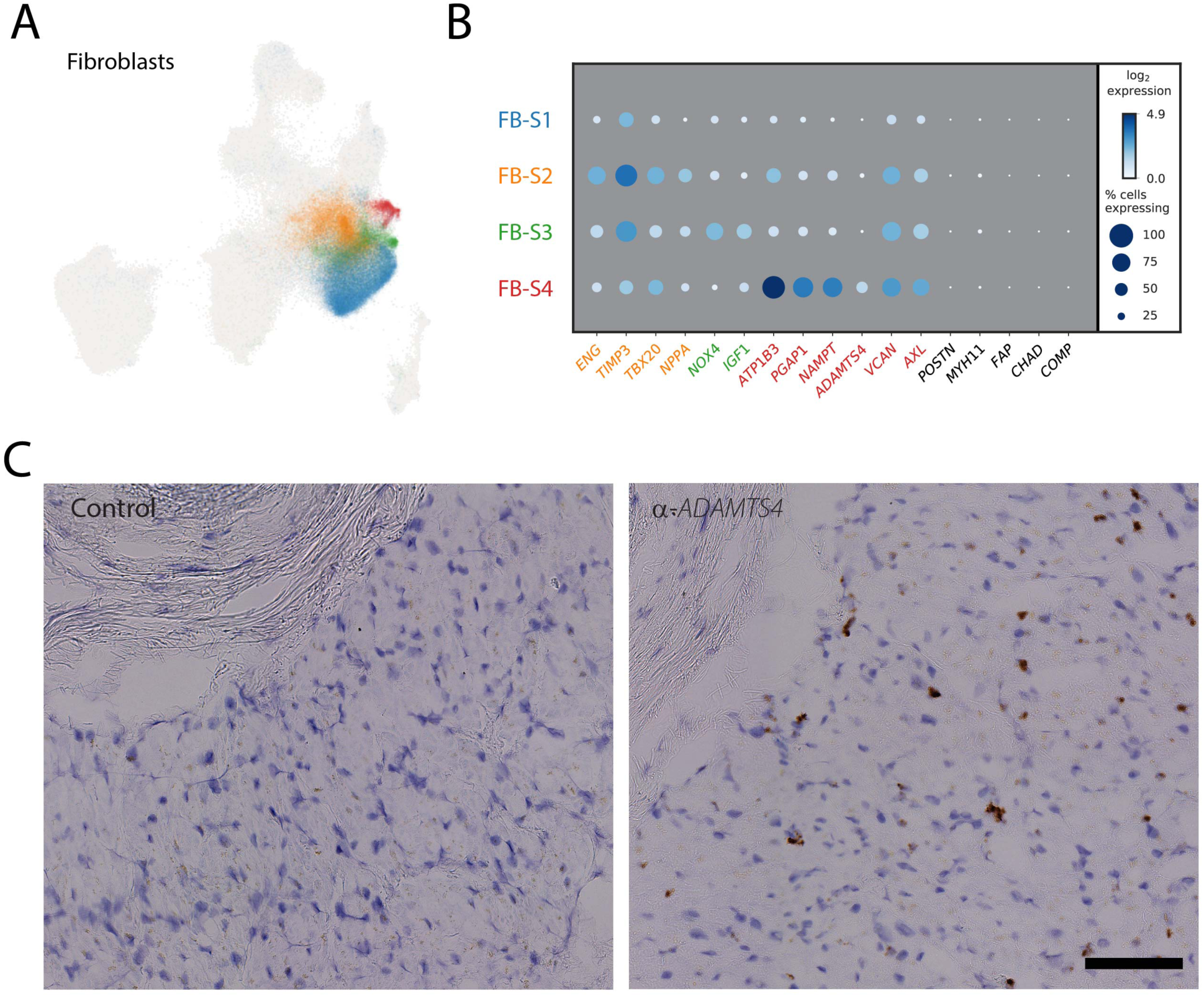
Subclustering fibroblasts to identify activated and quiescent fibroblasts. A: UMAP plot representing the four observed fibroblast subclusters superimposed over the global UMAP distribution. Each dot represents an individual cell and are colored by their respective subcluster B: Dot plot detailing the percentage of cells where each gene is detected (dot size) and mean log_2_ expression (blue hue) for representative subcluster marker genes. Each row represents the cell subcluster as displayed in panel A as according to color. C: Representative RNA *in situ* hybridization showing localization of *ADAMTS4* positive cells (brown stain) in sample LV1723 compared to a non-specific RNA probe (Control). Localization of nuclei is shown with hematoxylin (blue stain). Scale bar represents 100um.

#### Vascular support network of pericytes and vascular smooth muscle

Defining specific markers for microvessel associated pericytes and large vessel associated vascular smooth muscle cells has remained difficult, because the cells derive from similar progenitors and serve similar vascular support functions. We observed a relative enrichment of pericyte-specific *PDGFRB* in cluster 7 (AUC_7_=0.75) and the expression of smooth muscle actin (*MYH11*) in cluster 13 (AUC_13_=0.89) (Figure 2, **Table ST2**). This observation, combined with the preponderance of small vessels in our tissue samples, led us to classify the more numerous cluster 7 as pericytes and cluster 13 as vascular smooth muscle. Subcluster analyses of these cell types yielded little appreciable structure (**Figure S3B, Table ST3**), with the exception of cluster P-S2 in pericytes, which is enriched for some markers of endothelial cells (*VWF*, AUC_P-S2_=0.77, for example). Whether this indicates a differentiation event, as pericytes derive from endothelial cells, potential nuclear doublets, or ambient RNA contamination within the data, remains unclear.

#### A complex cardiac immune cell component

Two cell clusters (8 and 17) identified in this analysis have genetic signatures consistent with immune cell types. The first, cluster 8, represent cardiac resident macrophages and can be characterized by expression of the scavenger receptors *CD163* (AUC_8_=0.84) and *COLEC12* (AUC_8_=0.72), the mannose receptor *MRC1* (AUC_8_=0.85), the E3 ubiquitin ligase *MARCH1* (AUC_8_=0.72) and natural resistance-associated macrophage protein 1 (*NRAMP1* or *SLC11A1)* (AUC_8_=0.74) (**Table ST2**). Subclustering further revealed two populations that both express M2-polarization associated genes, including *RBPJ* and *F13A1* in M-S1 (AUC_M-S1_=0.85 and 0.84, respectively) and the transmembrane collagen *COL23A1* in M-S2 (AUC_M-S2_=0.65) (Figure 4A, **Table ST3**).

**Figure 4:**
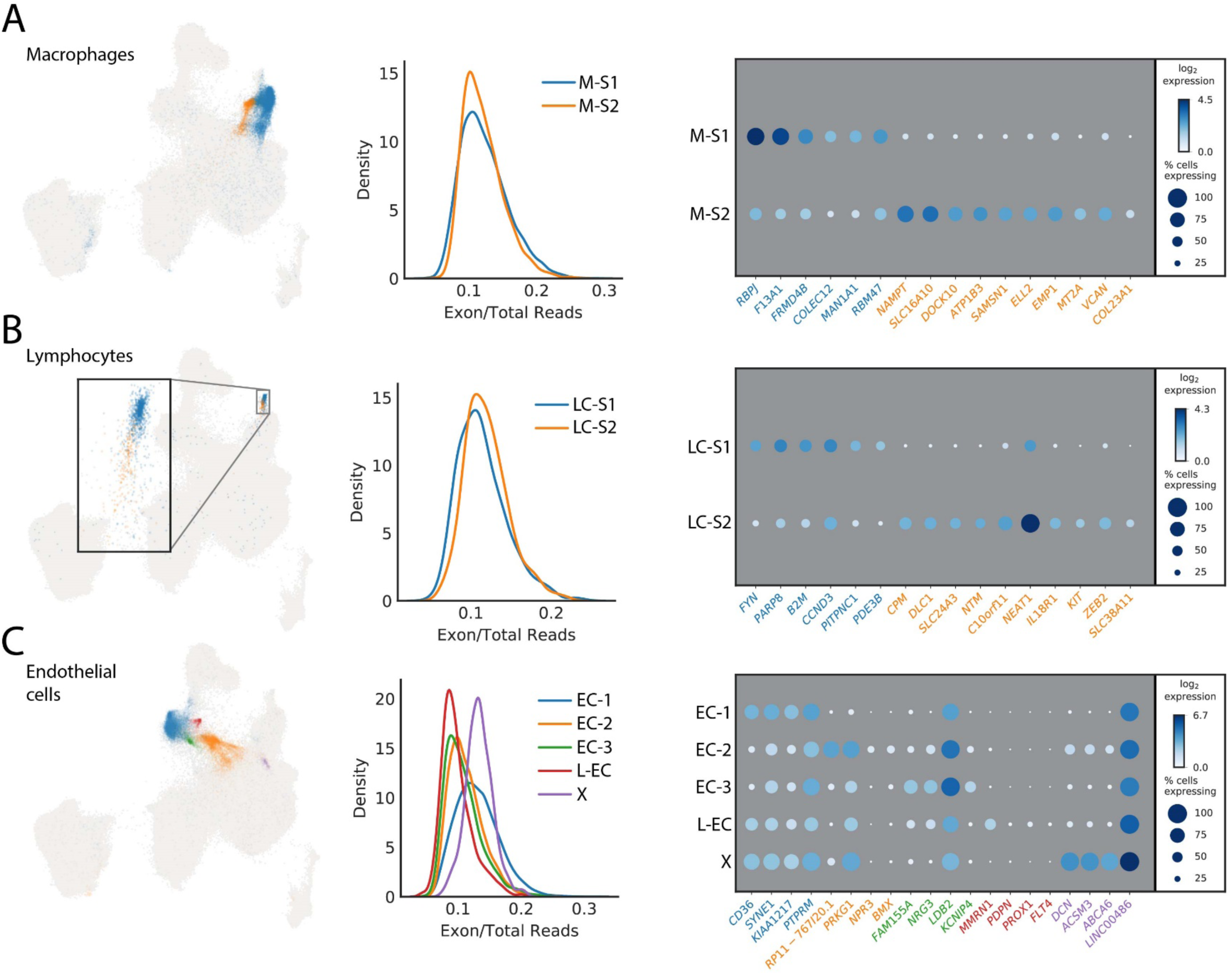
Subclustering to identify additional cellular diversity within macrophages, endothelial cells and lymphocytes. A: Left panel showing the UMAP distribution of the two identified macrophage subclusters. Each dot represents an individual cell colored by its respective subcluster. Center panel represents the calculated proportion of exonic mapping reads for the two subclusters. Right panel details the top markers by AUC for each subcluster. The size of the dot relates to the percentage of cells within the cluster which express that markers whereas the gradation relates to the mean log_2_ of the counts normalized by total counts per cell times 10,000. B: Left panel is the distribution of the two subclusters for lymphocytes in the global UMAP. Each dot represents an individual cell colored by its respective subcluster. Inset is the magnification of the outlined region. Center panel displays equivalent exon mapping reads for each of the subclusters. Right panel displays the top genes defining each subcluster as defined by AUC. C: Left panel is the distribution of the five identified subclusters of endothelial cells within the global UMAP plot. Each dot represents an individual cell colored by its respective subcluster. Center details the percentage of exon mapping reads, where cluster X (purple) has enrichment for exonic reads. Right panel shows a dot plot of the top markers for each subcluster by AUC with the addition of those markers used for identification of the lymphatic endothelium cluster derived from the standardized positive predictive value.

A second immune cell population (cluster 17) selectively expresses a number of well-known T cell markers. This includes the T cell surface antigen *CD2* (PPV50_17_=0.99), the early T cell activation antigen *CD69* (PPV50_17_=0.99), the T cell receptor associated transmembrane adaptor 1 (*TRAT1*) (PPV50_17_=0.98) (**Table ST2**). In addition, *PTPRC/CD45* (AUC_17_=0.77), an essential regulator of T- and B-cell antigen receptor signaling, the T cell immune adaptor *SKAP1* (AUC_17_=0.77), and the thymocyte selection marker *CD53* (PPV50_17_=0.91) show selectivity to this cluster (Figure 2, **Table ST2**). This overall lymphocyte population can be further subdivided into two distinct subclusters (LC-S1 and LC-S2). While many of the genes defining these subclusters have yet to be assigned functional roles within specific T cell populations, functional delineation may be enabled by future studies characterizing their localization within cardiac tissue and by analysis of transcriptional changes during disease. A gene of note within LC-S2 is *KIT*, which was long associated with cardiac resident stem cells, but since largely refuted [14]. We observe *KIT* expression exclusively within this lymphocyte subpopulation, with no evidence for expression in any cell with signatures of being progenitors or precursors for cardiomyocytes (Figure 4B and **Table ST3**).

#### Identification of vascular and non-vascular endothelial cells

The endothelial cell component of the heart consists of those cells which line the large and small circulatory vessels, the lymphatics, and the endocardium. From global clustering, we identified two major endothelial cell clusters (clusters 9 and 10), which express canonical markers such as *VWF* (AUC_9_=0.88, AUC_10_=0.77) and *PECAM-1* (AUC_9_=0.71, AUC_10_=0.81), but were unable to further resolve subtypes prior to subclustering analysis (Figure 2, **Table ST2**).

Five subclusters were identified within combined endothelial clusters 9 and 10 (Figure 4C). We were unable to clearly resolve subclusters based on AUC markers alone, but interrogation of less abundant genes with significant selectivity proved useful in identifying subcluster populations. For instance, in subcluster 4 (L-EC), we observed enrichment for cells expressing lymphatic endothelial cell markers including *PROX1*, *FLT4* and *PDPN* (PPV50_L-EC_ of 0.95, 0.91, and 0.94, respectively) (**Table ST3**). A subset of cells in EC-S2 express *BMX* (AUC_EC-S2_=0.65), an artery specific endothelial cell marker as well as *NPR3* (AUC_EC-S2_=0.65). In mice, *NPR3* is selectively expressed in adult endocardium [15], suggesting the EC-S2 population may represent endocardial cells (**Table ST3**). These observations reflect the fact that the heart biopsies used did not include any large vessels, explaining in part the lack of distinct arterial and venous endothelial cell populations.

#### Epicardial adipocytes enriched in the leukocyte marker CD96

Epicardial adipose tissue is present in human hearts which comprises up to 20% of its total mass.[16] Adipocytes may also be observed within the heart itself in pathological conditions such as obesity or cardiomyopathy. Tissues were generally free of myocardial adiposity as observed by histology in our samples with the exception of the right ventricle of P1723 (**Figure S4C**). Given that cells of this sample are not overly represented in the cluster, we propose that Cluster 11 is comprised primarily of epicardial adipocytes, with ontology analysis identifying terms such as fatty acid and lipid metabolism (Figure 2).

These cells were characterized by genes whose expression ultimately regulate the size and stability of lipid droplets, such as *CIDEC* (AUC_11_=0.72) and *PLIN5* (AUC_11_=0.78). These data also support the view of epicardial fat as an endocrine organ. *ADIPOQ*, which modulates fatty acid transport and increases intracellular calcium is present in nearly 65% of adipocyte nuclei but only 0.3% of other cell types (AUC_11_=0.82). Within this population, *TRHDE*, which inactivates thyrotropin releasing hormone, and *IGF-1* are also strongly enriched within this population (AUC_11_=0.76 and AUC_11_=0.76, respectively). *IGF-1* also has an important role in cell growth, proliferation and resistance to death later in an individual’s life, functions which directly relate to its significant role in the development of obesity.[17] Surprisingly these cells are also enriched for *CD96*, a marker most often identified with Natural Killer (NK) and T-cells (AUC_11_=0.73) (**Table ST2**).

#### Autonomic neuronal inputs of the intrinsic cardiac network

The heart is innervated by the central nervous system through the cardiac plexus, which distributes parasympathetic (vagal) and sympathetic stimulation. In addition, an intrinsic cardiac autonomic network, consisting of ganglionated plexi within epicardial fat pads, resides within all four chambers of the heart. We identified a subset of neuronal cells in cluster 16, largely defined by neuronal cell adhesion genes such as the neurexins (*NRXN1,* AUC_16_=0.91 and *NRXN3,* AUC_16_=0.87), and *NCAM2* (AUC_16_=0.73) rather than by electrophysiology or secretory associated genes. The only ion channel gene identified as a marker in this cluster is *SCN7A* (AUC_16_=0.74), initially described in glia, but since understood to reside in other cell types of the nervous system [18]. For signaling genes, the receptor genes *ADGRB3* (AUC_16_=0.72), which acts to promote angiogenesis, and *SHISA9* (AUC_16_=0.72), which modulates AMPA-type glutamate receptors, were robustly expressed within this cluster (**Table ST2**). Given the sampling location of the lateral wall and the presence of this neuronal subtype through all four chambers, it is likely that the neuronal cells identified within the present study are derived from the intrinsic cardiac autonomic network.

### Differential expression analysis uncovers chamber- and sex-specific gene expression profiles within cell subtypes

We next determined whether expression programs in the major cell types differed by cardiac chamber or sex. Prior to performing differential expression testing, we first removed any cluster or subcluster that was previously labeled as cytoplasmic (clusters 5 and 12 from the global map and subclusters CM-S5 and EC-S5), collapsed cell clusters into their respective major cell types, and removed genes with a poor PPV50 for the major cell type of interest (**Methods**). We then performed differential expression testing using a generalized linear mixed model framework on the 5 most numerous major cell types (cardiomyocytes, fibroblasts, endothelial cells, pericytes, and macrophages). The smaller number of cells for other cell types coupled with the sparsity of single nucleus RNA-sequencing expression matrices sequencing limited our ability to confidently call differentially expressed genes in rare cell types.

#### Cardiomyocytes are the most distinct cell type between chambers

Atrial and ventricle cardiomyocytes are well known to have distinct physiological functions, contractile properties, and electrical signaling. These functional and structural differences are reflected in discrete transcriptional profiles. As anticipated, when we compared the atria to the ventricles we observed a total of 2,300 genes that reach an FDR adjusted significance threshold (Figure 5A,B,C **Table ST4**). These differences were exemplified by an increased expression of *HEY2* and *MYH7* in the ventricles (effect size=3.75, *P*=1.5×10^-27^ and effect size=1.83, *P*=1.07×10^-16^, respectively), and *NPPA* and *MYL4* in the atria (effect size=6.89, *P*=1.22×10^-30^ and effect size=4.23, *P*=2.94×10^-29^, respectively). We identified 2,058 differentially expressed genes between the left atrium and left ventricle, but only 1,134 differentially expressed genes between the right atrium and right ventricle.

**Figure 5:**
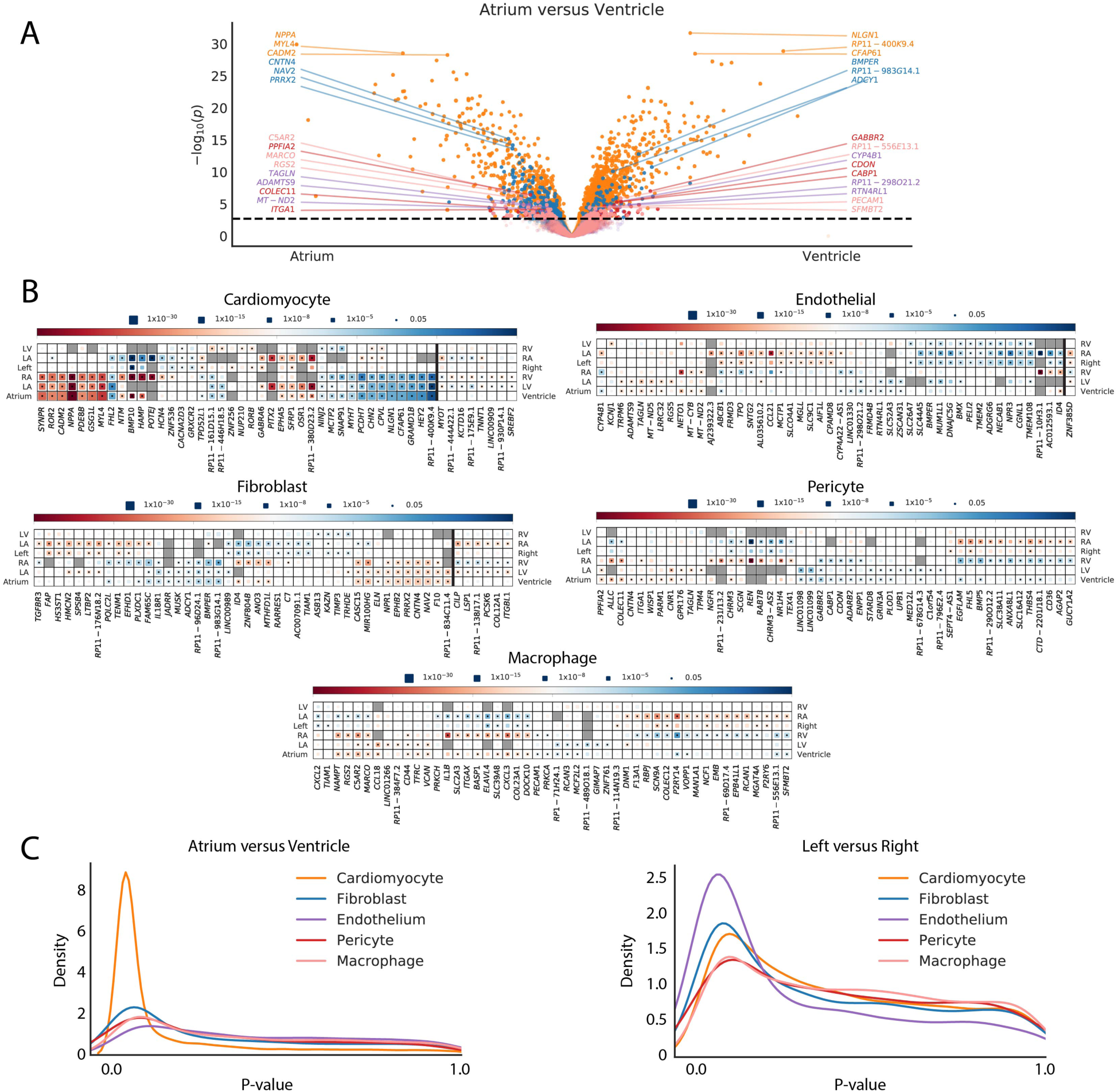
Differential expression analyses for chamber specific signatures of major cell types. A: Volcano plot detailing differential expression of genes when comparing the aggregated atrial and ventricular chambers in cardiomyocytes (orange), fibroblasts (blue), endothelial cells (purple), pericytes (red), and macrophages (pink). The X-axis represents the fixed effect from the generalized linear mixed model and the Y-axis represents the -log_10_(P-value). Dotted line indicates the FDR adjusted P-value threshold for statistical significance. The top 3 genes upregulated in atrial cells and ventricle cells are highlighted for each cell major cell type. B: Heat maps detailing a representative selection of significantly differentially expressed genes between chambers within major cell types. Color indicates whether the gene is enriched within the chamber listed on the left (red) or right (blue). Size of the inset block indicates the P-value for the comparison. Dot within the block indicates statistical significance for the given comparison. Genes to the right of the dark vertical line are those with different directionalities when comparing atria versus ventricles on the left or right side. C: Density plot displaying the number of genes with certain P-values across the P-value spectrum within each major cell type for atrium versus ventricle (left panel) and left versus right (right panel) comparisons.

In contrast to the marked transcriptional patterns observed between the atria and ventricles, there were many fewer genes that were differentially expressed when comparing the left versus the right side of the heart. A comparison of the left versus right atria revealed 248 differentially expressed genes, while only 24 genes were differentially expressed between the left and right ventricles.

Closer inspection of the data yield noteworthy insights into chamber specific expression programs. For example, the atrial fibrillation susceptibility gene, *PITX2* [19] was observed in 2.3% of left atrial cardiomyocytes and in less than 0.05% of cardiomyocytes in any other chamber. Interestingly, *HCN4* is present in 4.3% of cardiomyocytes from the right atrium, in only ∼1% of cardiomyocytes from the right ventricle and left ventricle, and less than 0.5% of cardiomyocytes from the left atrium. The *HCN4* gene encodes the ion channel responsible for spontaneous depolarization and has also been associated with atrial fibrillation.

Other genes with limited or entirely unexplored roles in cardiomyocyte biology also exhibit chamber preference. Among these, *HAMP*, which encodes a protein for regulating iron export, and the solute carrier gene *SLC5A12* are found predominantly within the right atrium (present in 18.3% and 5.8% of cardiomyocytes in the right atrium, respectively, compared to < 1% of cardiomyocytes in any other chamber). Eight genes display significant differences in expression in opposing directions when comparing left or right atrium to their respective ventricular partner (Figure 5B**, Table ST4**). Among these are *MYOT* (left: effect size=0.75, *P*=8.86×10^-5^; right: effect size=-0.93, *P*= 2.00×10^-5^) and *TNNT1* (left: effect size=0.64, *P*=0.001; right: effect size=-0.45, *P*=1×10^-4^), which are enriched in the right ventricle and left atrium and which play critical roles in sarcomeric organization and function.

#### Non-cardiomyocytes display striking chamber-specificity

While differences in cardiomyocytes between chambers were expected, it was less clear from previous work if chamber specificity exists within other cardiac resident cells. Surprisingly, there were profound signatures of chamber specificity in the other cell types examined. A total of 765 genes surpassed FDR-corrected *P*-value threshold in fibroblasts for at least one comparison of chamber or laterality. In addition, 125 genes in pericytes, 320 genes in macrophages and 354 genes in endothelial cells were also found to be differentially expressed. (Figure 5B**, Table ST4**).

Among fibroblasts, pericytes and macrophages, the atrial versus ventricular comparisons account for the majority of differential expression, with the right atrial cells being consistently the most divergent. In some cases, this divergence is sufficient to drive some of the subclustering observed within Figure 4 and **Figure S3**. The most striking example of this is within the macrophage population, where the differential expression between the right atrial macrophages and those of other chambers is strong enough to detect a second macrophage subcluster (M-S2, Figure 4A) which consists almost entirely of right atrial cells (94.0%). In contrast, endothelial cells are most distinct when comparing sidedness (220 differentially expressed genes genes for left versus right, 43 differentially expressed genes for atrium versus ventricle). Again, much of this is driven by the right atrium, with 217 significant genes when comparing to the left atrium. This difference manifests within the subclustering, where right atrial cells make up 88.2% of subcluster EC-3 (Figure 4C).

Similar to the cardiomyocytes, some genes display different directionalities when comparing atria versus ventricles on the left or right side (Figure 5B**, Table ST4**). This includes 6 genes in fibroblasts, including *CILP* (left: effect size=1.13, *P*=4.04×10^-5^; right: effect size=-1.55, *P*=8.43×10^-7^) and *ITGBL1* (left: effect size=1.07; *P*=7.26×10^-6^; right: effect size=-0.67; *P*= 4.08×10^-4^) which have links to the regulation of fibrosis [20, 21] and 1 gene in endothelial cells, *ZNF385D* (left: effect size=0.82, *P*=0.001; right: effect size=-1.14, *P*=1.60×10^-8^). In sum, there are profound differences in the expression profiles of non-myocytes across the cardiac chambers.

#### Sex-based differential expression identifies genes associated with myopathy and coronary artery disease

Biological sex has profound impact upon cardiac morphology, physiology, and susceptibility to cardiovascular disease, but the molecular differences of the heart between the sexes remain obscure. Given the inclusion of 4 female and 3 male donors within our data, we proceeded to separate the cells by sex and performed differential expression testing within the same 5 major cell types both globally and by chamber of origin. Given our limited sample size, the number of sex-specific genes was greatly reduced when compared to those derived from chamber specificity in the previous section. In total, 17 genes exhibited sex-based differential expression within cardiomyocytes, 2 within the endothelium, 10 within the fibroblasts, 3 within the macrophages, and none for the pericyte comparisons (**Table ST5**). Approximately one third of the genes that were differentially expressed by sex were autosomal (Cardiomyocyte = 6, fibroblast = 4). An anticipated, several of these differentially expressed genes are related to hormonal signaling. *CRISPLD2* is induced by the progesterone receptor [22] and *UGT2B4* is involved in estrogen metabolite modification.[23] *NEB*, which encodes the sarcomeric structural protein nebulin, is enriched within the left ventricle in males (effect size=1.54, *P*=1.73×10^-6^) while *ZNF827*, which resides proximal to a GWAS locus for coronary artery disease [24] is expressed at increased levels in women with the most marked upregulation in the right atrium (effect size=1.31, *P*=2.12×10^-6^).

#### Integration of single nucleus RNA-seq data with cardiovascular genetics and the druggable genome

We next sought to apply our snRNA-seq data to better understand the basis of human cardiovascular disease using three complementary approaches. First, we examined the cell type specific expression of genes implicated in Mendelian forms of cardiovascular disease. Second, we related cardiac transcriptional data to the data derived from population-based, genome wide association studies (GWAS) for cardiovascular diseases and traits. Finally, we intersected our snRNA-seq data with genes that are potentially druggable in order to identify novel therapeutic targets for cardiovascular diseases.

#### Genes implicated in cardiomyopathies and arrhythmia syndromes are enriched in cardiomyocytes

Intersection of our snRNA-seq data with a panel of genes previously implicated in cardiomyopathies and arrhythmia syndromes revealed three 3 general patterns. First, as anticipated, over 25% of the pathogenic genes show enriched selectivity (AUC > 0.70) in the cardiomyocyte population (Figure 6**, S5**, 17/75 genes for arrhythmias (p < 0.0001), 27/106 genes for cardiomyopathies (p < 0.0001)). Second, a smaller subset of known pathogenic genes are highly expressed in non-cardiomyocyte populations. This pattern was exemplified by the *ABCC9* gene which has been implicated in dilated cardiomyopathy and is predominantly expressed in pericytes. Similarly, *LAMA4*, which encodes a component of the extracellular matrix and has been associated with dilated cardiomyopathy, was specifically expressed in adipocytes (AUC_AD_=0.79). Finally, we found that approximately half of the genes implicated in Mendelian cardiovascular diseases were not highly or broadly expressed in the healthy human heart.

**Figure 6:**
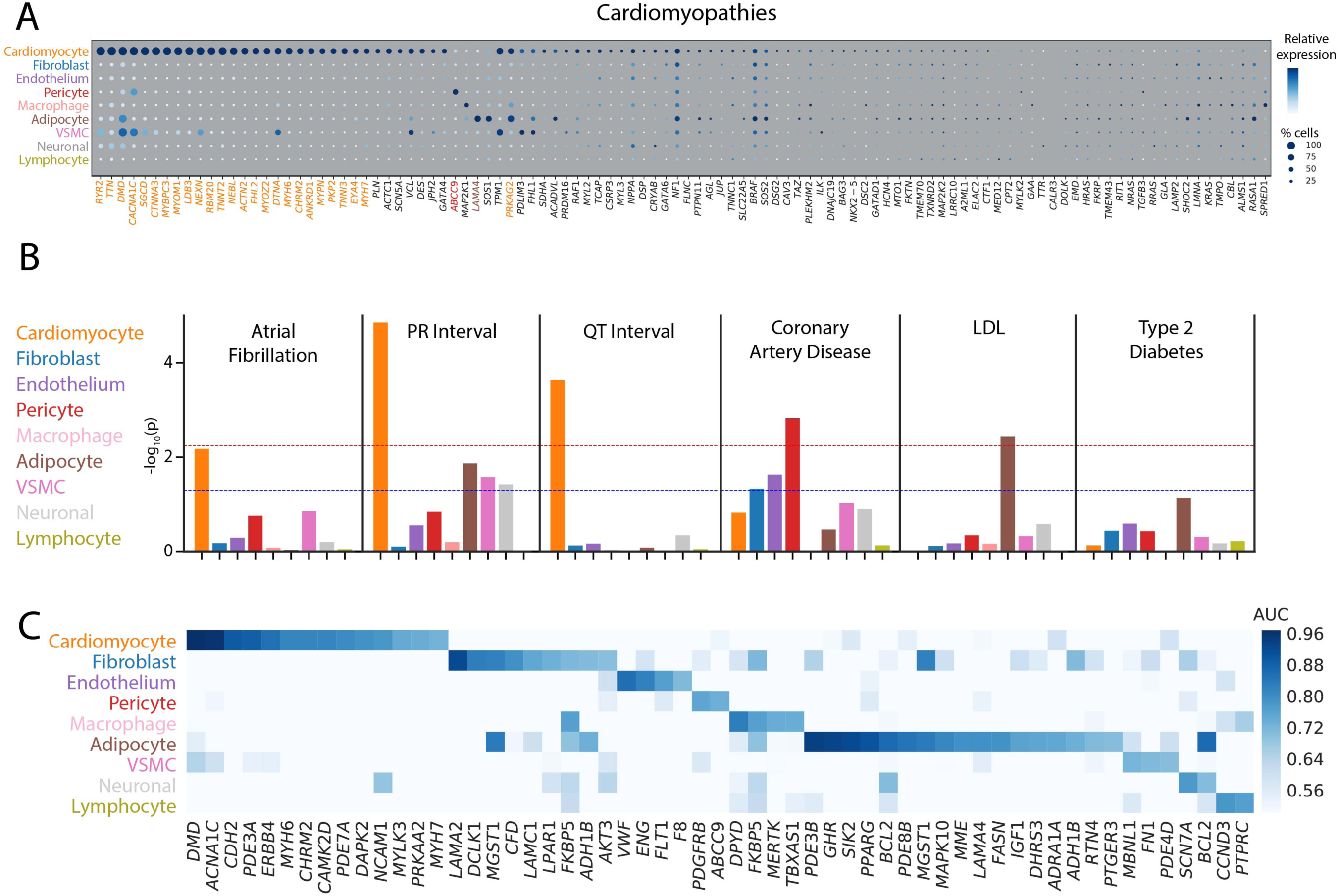
Integration of single nucleus RNA sequencing with genetic associations to uncover disease biology. A: Dot plot for genes currently on standard cardiomyopathy clinical testing panels. The size of each dot represents the percent of cells in which the gene of interest is detected and the shading represents the relative expression of the gene. Color of the genes correspond to the cell type for which the AUC reaches 0.70 or greater. Genes with black color indicate no cell type which reaches this threshold. Size and shade of the dot corresponds percentage of cells and relative expression, respectively. B: Results of LD score regression analyses on the combined major cell types. Dotted lines display unadjusted (blue) and Bonferroni adjusted (red) P-value thresholds for statistical significance. Colors of the bars correspond to the color of the cell major cell type labels on the left. C: Heat map detailing the intersection between single nucleus RNA sequencing data and Tier 1 druggable genes. Genes with an AUC greater than 0.70 in at least one cell type are shown. Shade of the color represents the AUC value for the gene within each cell type. Of note, genes that have an AUC greater than 0.70 in multiple cell types appear multiple times in the plot.

#### Combining GWAS and snRNA-seq data to identify the most relevant cell types for cardiovascular diseases

To identify putative cell types of interest to a set of complex traits and diseases, we employed linkage disequilibrium (LD) score regression to partition genetic heritability from GWAS studies. Briefly, assuming a cis-regulatory model for single nucleotide polymorphism (SNP) function, the approach partitions SNP heritability derived from GWAS across regions near genes considered to be cell type specific in our sn-RNAseq data. Should SNP-trait associations be enriched around cell type specific genes, this suggests that heritability of the trait is driven in part by the genetic effects in that cell type. We applied this approach to a range of cardiometabolic traits, as shown In Figure 6B.

Integration of our single nucleus sequencing results with GWAS data for cardiometabolic traits revealed the expected enrichment in cardiomyocytes for two electrocardiographic traits, the PR interval (*P*=1.4×10^-5^) and the QT interval (*P*=2.3×10^-4^). We observed a similar cardiomyocyte enrichment for the most common cardiac arrhythmia, atrial fibrillation (*P*=0.007). Interestingly, we also observed a marked enrichment in pericytes for genes at the loci for myocardial infarction (*P*=0.001) and in adipocytes for LDL cholesterol (*P*=0.004).

After examining global enrichments, we chose to employ a more reductionist approach to evaluate potentially unique expression profiles of disease-associated genes. Expression quantitative trait loci (eQTL) mapping, which evaluates changes in gene expression due to genotype, is a common strategy for linking a GWAS locus to a particular gene. We used the intersection of known disease or trait associated eQTLs from GTEx [25] and our own work [26] to determine the cell type where the transcript of interest is most highly expressed. For each trait, we limited our analysis to genes from the most disease relevant tissue, for example the QT interval is only intersected with left ventricular eQTLs and atrial fibrillation only those from the left atrium. eQTLs are derived from tissue level RNAseq experiments, and are thus predisposed to discover signals in more prevalent cell types. Surprisingly, rather than patterns which indicate cardiomyocyte centered expression, genes generally show non-specific cell type expression, with a few interesting patterns emerging (**Figure S5B**). Within the left ventricle, 1 of the 11 putative genes for PR interval (*PDZRN3*) shows enriched expression in cardiomyocytes (AUC_CM_=0.88), 1 of 21 putative genes for QT interval (*SLC35F1*) shows enriched expression in neuronal cells (AUC_NR_=0.71), and 2 of 37 putative genes for CAD show enriched expression in adipocytes (*C6orf106*, AUC_AD_=0.70) and vascular smooth muscle cells (*LMOD1*, AUC_VSMC_=0.74). Interestingly, in the left atrium, the putative PR interval gene *PDZRN3* shows enriched expression in adipocytes (AUC_AD_=0.76) and 2 of 12 atrial fibrillation genes show enriched expression in cardiomyocytes (*CASQ2*, AUC_CM_=0.74) and endothelial cells (*SYNE2*, AUC_EN_=0.74).

#### Cell-type specific expression of potentially druggable genes

To identify potential drug targets in cardiac tissue, we sought to identify tier 1 classified genes from the druggable genome [27] that shows selectivity toward particular cardiac cell types. This tier includes targets of both approved drugs and those in clinical development. Of the 1420 potential genes, 53 unique genes were specifically expressed in at least one major cell type with an AUC > 0.70 (Figure 6A). Most commonly these genes were found in adipocytes (n=17), cardiomyocytes (n=14), and fibroblasts (n=9). Among these, *CACNA1C*, the receptor for calcium channel blockers that are commonly used to treat hypertension, and *PDE3A,* a known target of inamrinone for treatment of congestive heart failure,[28, 29] showed selectivity toward cardiomyocytes. However, the selective expression of other druggable genes in cardiac cell types, and particularly in non-myocytes, will provide new opportunities for future therapeutic development.

## Discussion

We have developed a comprehensive map of the transcriptional landscape in normal human heart comprised of snRNA-seq for more than 280,000 cells. Our work provides at least four novel advances that will enhance our understanding of cardiovascular biology. First, we have developed the largest collection of single nuclear transcriptomes from the human heart to date. This robust dataset allowed us to define 9 major clusters and at least 20 subclusters of cell types within the healthy heart. Second, we identified unexpected differences in chamber-, laterality-, and sex-specific transcriptional programs across major subtypes of cardiac cells. Third, we linked specific cell types to common and rare genetic variants underlying cardiovascular diseases. Finally, we generated a portable, scalable analytic and statistical framework for handling the unique challenges of cardiac single nuclear data that will be of broad interest to the scientific community.

Previous single cell sequencing of the heart has focused on murine models of health and disease [4,7,30–33], with limited forays into analyses of human tissues [5, 6]. Notable examples of the latter include compelling studies of fetal development and cardiomyopathy-control comparisons. The rarity of data from humans highlights the inherent technical and logistical challenges associated with these studies. Ideal tissue harvesting requires coordination between clinical and laboratory teams to quickly isolate and preserve the metabolically active, ischemia-sensitive tissue. After tissue isolation, additional challenges emerge, including problematic cell isolation protocols combined with large disparities in cell size necessitating nuclear rather than whole cell sequencing. Further, the lysis of cells for single nuclear isolation produces significant cytoplasmic RNA contamination in the form of ambient RNA, which we remove using a probabilistic model developed by our group. In human tissue, there is also significant intersample diversity such that cell alignment across samples is required for any additional cell subtype comparisons. As batch correction with the commonly used canonical correlation analysis (CCA) may remove sample specific clusters [34], we applied a deep neural network to correct batch effects using the scVI tool [35]. Finally, the transcriptional complexity of nuclei is not equivalent between cell types, making identification of droplets containing cells versus those which are empty more challenging than typical cell-based protocols. To overcome this challenge, we called cells using our CellBender *remove-background* tool which compares each droplet to the background signature of ambient RNA to identify and retain cell types with lower average transcriptional coverage.

The result of highly collaborative effort is a large-scale map of the transcriptional diversity of the human heart that is approximately 50 times larger than prior human studies. The scope of our study afforded us the ability to interrogate rarer cell types, perform detailed cellular subclustering, and define the signatures of cell types beyond what was previously possible. We believe that our data will be a unique resource for the cardiovascular research community and is available for further exploration at the Broad Institute’s Single Cell Portal (https://portals.broadinstitute.org/single_cell). This data will facilitate the independent evaluation of the cell types we have described, provide the opportunity for re-analyses and more liberal cellular subclustering, examination of the expression of genes of interest, and additional comparisons across and within cell groups.

Beyond analyses we have presented here, we anticipate that this work will serve as a framework for further studies, both as a reference dataset of human non-failing samples, and as an analytic framework for further comparisons. We were excited to read the initial studies of human disease comparisons by single cell sequencing, and hope that the data and approach here will facilitate further comparisons of this kind in the future. As highlighted with the discovery of ionocytes based on CFTR expression in patients with cystic fibrosis [36], we hope to identify similar rare disease-specific cellular subtypes that can be used in cardiovascular disease research. Looking forward, recent advances in the non-cardiovascular single cell work using LIGER [37] and Seurat v3.0 [38] have highlighted the potential for multi-modal integration of transcriptome and epigenome datasets. Generation of richer datasets of this nature in these samples and others will further facilitate translational discoveries, while overcoming limitations of any single data modality. Finally, we hope that this is the first entry in a larger series of large human transcriptomes to be published by our group and others. When combined, these data can facilitate analyses which require significant sample sizes, such as eQTL analyses which link risk loci to genes; these methods are just beginning to be applied to single cell data [39].

### Limitations

Our study was subject to several potential limitations. Although this is a much larger collection of human cardiac transcriptomes than any other study to date, these individuals may not reflect the complete diversity contained within non-failing hearts. Studies to expand the number of normal and diseased tissue comparisons are ongoing. Second, all individuals in this study were of European descent; thus, transcriptional profiling of samples from other races and ethnicities should be a goal in the future. Third, sex-based comparisons were relatively underpowered given the limited numbers of individuals present in the study. Fourth, nuclear transcriptomes represent a small percentage of the total mRNA present in a cell and differ significantly from the population of species present in the cytoplasm. Follow up studies that examine the concordance of whole cell versus nuclear transcriptomes will clarify the differences in these two populations of mRNA. Finally, methods to remove ambient RNA, identify nuclear doublets, perform batch correction are imperfect; even after correction droplets are expected to retain some background signal. Thus, interpretation of the data should keep this in mind, especially when observing the expression of genes from common cell types, such as cardiomyocytes, in other cell types.

### Conclusions

Single cell RNA sequencing has been a revolutionary tool for characterizing known and novel cell types and states in health and disease. Here we provide a large-scale map of the transcriptional and cellular diversity in the normal human heart. Our identification of discrete cell subtypes and differentially expressed genes within the heart will ultimately facilitate the development of new therapeutics for cardiovascular diseases.

## Author Contributions

Conceptualization: NRT, MC, CS, PTE

Methodology: NRT, MC, SJF, AWH, VAP, KB, ADA, CNH, AA, FA, MB, KBM

Software: SJF, MB, FA

Validation: NRT, VAP, CNH

Formal Analysis: MC, SJF, AWH, MB

Investigation: NRT, CNH, VAP

Resources: FA, KGA

Data Curation: MC, KB, CR, KBM

Writing – Original draft: NRT, MC, PTE

Writing – Review and Editing: SJF, AWH, VAP, KB, ADA, CNH, AA, IP, CR, FA, SHC, KGA, MB, KBM, CS

Visualization: NRT, MC, SJF

Supervision: NRT, PTE

Project Administration: CS, PTE

Funding Acquisition: CS, PTE

## Methods

### Human tissue samples

Adult human myocardial samples of European ancestry were collected from deceased organ donors by the Myocardial Applied Genetics Network (MAGNet; www.med.upenn.edu/magnet). For all donors, clinical examination and medical history displayed no indications of structural heart disease. Employing methods used in clinical transplantation, all hearts were arrested in situ with at least 1 liter of ice-cold crystalloid cardioplegia solution, as previously reported.[40, 41]Hearts were transported to the lab in ice-cold cardioplegia solution until cryopreservation (always <4 hours). Written informed consent for research use of donated tissue was obtained from next of kin in call cases. Research use of tissues were approved by the relevant institutional review boards at the Gift-of-Life Donor Program, the University of Pennsylvania, Massachusetts General Hospital and the Broad Institute.

### Single nucleus RNA-sequencing

Single nucleus suspensions were generated by a series of cellular membrane lysis, differential centrifugation and filtration steps. Approximately 100mg of tissue was cryosectioned at 100 um on a Leica CM1950 cryostat to enable liberation of nuclei from the tissue while minimizing mechanical manipulation. Tissue sections were homogenized in a dounce homogenizer after suspension in 4mL of ice cold lysis buffer containing propidium iodide for nuclear staining (250mM Sucrose, 25mM KCl, 0.05% IGEPAL-630, 3mM MgCl2, 1uM DTT, 10mM Tris pH 8.0). After 5 minutes incubation, large debris was pelleted at 20g for 1 min in a Beckman Coulter Allegra X-15R swinging bucket centrifuge. Supernatant was brought to 8mL of total volume with nuclear wash buffer (PBS + 3mM MgCl2 + 0.01% BSA) then filtered sequentially through a 100um and 20 um filter (pluriSelect Life Science). Nuclei were pelleted at 400g for 5 minutes at 4C, washed in 4mL of nuclear wash buffer and repelleted. After removal of wash buffer, nuclei were resuspended in approximately 500uL of cold nuclear resuspension buffer (Nuclear wash buffer + 0.4U/uL of murine RNAse inhibitor (New England Biolabs)) with gentle trituration then counted on a hemocytometer. Cells were loaded into the 10x Genomics microfluidic platform (Single cell 3’ solution, v2) for an estimated recovery of 5000 cells per device. Processing of libraries was performed according to manufacturer’s instructions with a few modifications. First, nuclei were incubated at 4C for 30 minutes after emulsion generation to promote nuclear lysis. Second, the reverse transcription protocol was modified for one of the two replicates to be 42C for 20 minutes then 53C for 120 minutes. This is noted as (_2) in the Sample ID column of the sample information table (Supplemental Table ST1). Libraries were multiplexed at an average of 4 libraries per flow cell on an Illumina Nextseq550 in the Broad Institute’s Genomics Platform.

### Sample selection and quality control

In total, 56 single nuclei RNA-seq experiments were performed from all four chambers of the human heart in 7 distinct biological individuals, processed in duplicate. Reads from single nuclei experiments were de-multiplexed and aligned to a GRCh38 human pre-mRNA reference using the 10x Genomics toolkit CellRanger 2.1.1 and default parameters with the exception of setting the --expect-cells flag to 5000 based on library preparation.

For each experiment, the distribution of the number of unique molecular identifiers (UMI) was visually inspected to identify experiment failures. Any experiment where the fraction of reads in cells was less than 33% (n=7) or the median UMI per cell was less than 600 (n=6) based on CellRanger were excluded from further analysis. In total 9 experiments failed on these criteria. These cutoffs corresponded to poor structure in the UMI decay curve.

Additionally, genetic concordance was checked between all experiments of the same biological individual using the Genome Analysis Toolkit (GATK) [42] method *CrosscheckFingerprints* on the single cell 10x aligned reads. A list of approximately 6,300 sites was provided and samples were considered concordant if their corresponding LOD score was greater than 10. Two experiments from the right ventricle of individual **P1681** were discordant with the remaining experiments from the same individual and were subsequently removed.

### Post-Sample Selection Processing: CellBender

All 45 experiments passing initial quality control were processed using the *remove-background* tool from CellBender v0.1 to determine which droplets contain a cell and to correct gene count matrices by removing ambient background RNA contamination. For complete details on the CellBender *remove-background* model, see https://github.com/broadinstitute/CellBender. Briefly, CellBender performs Bayesian inference in the context of a probabilistic model to remove ambient RNA by estimating the contribution of ambient background RNA captured in each droplet and adjusting the count matrix appropriately. The CellBender model does require that there are some unambiguously empty droplets containing only ambient, background RNA. One additional experiment was removed because of poor definition in the UMI decay curve which prevented CellBender from converging appropriately.

Specifically, CellBender *remove-background* was run on a Tesla K80 GPU using the following parameters: expected-cells 5000, total-droplets-included 20000, low-count-threshold 50, epochs 300, z-dim 200, z-layers 1000, empty-drop-training-fraction 0.3. After an examination of the output cell calls, it was determined that unusually high ambient RNA had led to a failure on sample RV_1666_2, which was subsequently rerun with the following parameters altered: total-droplets-included 15000, low-count-threshold 200.

### Four chamber map aggregation

Prior to cell clustering, additional low quality cells were removed on a per-experiment basis to account for large variability in sequencing depth and complexity between experiments. These pre-processing steps were performed using Seurat 2.3.4. In brief, cells were removed from an experiment if: 1) the number of genes detected was less than 100 or greater than a predefined upper outlier cutoff, 2) the number of UMI for the cell was greater than a predefined upper outlier cutoff, or 3) the percent of mitochondrial gene content was greater than 5%. The upper outlier cutoff was calculated as the third quartile plus 1.5 times the interquartile range. Upper cutoffs were used to minimize the introduction of multiplets into downstream clustering. These criteria reduced the total number of putative cells from 373,243 to 287,269 for subsequent aggregation.

Highly variable genes were selected to perform cell clustering using Seurat 2.3.4. The aggregated cell count matrix was first normalized by dividing the number of UMI for each transcript by the total UMI for the cell, multiplying by 10,000, and taking the natural log of these results. Variable genes were found globally using the *FindVariableGenes* function in Seurat which bins genes by average expression and calculates a *Z* score for dispersion within each bin [1]. Normalized expression bounds were set between 0.03 and 5, and genes with dispersion *Z* score greater than 0.5 were selected. In total, 1,969 genes remained for clustering.

Because of large biological heterogeneity between samples, we applied the single-cell variational inference (scVI) framework to map cells from different samples onto a joint coordinate system.[35] In brief, scVI uses deep neural networks to learn the underlying distributions of cell-level expression, while accounting for batch variables. By treating the individual, rather than the experiment, as a batch indicator, this procedure aligns cells accounting for heterogeneity between individuals. Note that the success of scVI in removing biological batch effects implies that in our dataset, batch effect is dominated by biological inter-individual variability and is not technical in nature. The inferred latent space can then be used for downstream clustering of cells. We applied scVI 0.3.0 on the previously identified 1,969 genes to estimate 50 latent variables. A neighborhood graph of cells was built based on these 50 latent variables using scanpy 1.4 [43](*scanpy.pp.neighbors*) selecting a cosine distance metric and using 15 neighbors. Cells were subsequently placed into clusters using the Louvain algorithm (*scanpy.api.tl.louvain*) with default parameters and a resolution parameter of 1.0. To visualize cells in a high dimensional space, uniform manifold approximation and projection (UMAP) was applied to the same latent space using a cosine distance metric, and default parameters.[44]

### Calculation of intronic and exonic reads

scR-Invex (Aaron Graubert, François Aguet; https://github.com/broadinstitute/scrinvex) was run in order to count the reads in each BAM file output by CellRanger 2.1.1 *count* that mapped to intronic, exonic, and junction regions of the transcriptome.

### Sub-chamber visualization

Sub-chamber maps were generated using global latent variables and retaining cluster identities from the global cell map. Default UMAP parameters were used with the exception of setting metric=’cosine’, spread = 1, min_dist = .3, and n_neighbors = 25 for left ventricle and metric=’cosine’, spread = 1, min_dist = .1, and n_neighbors = 15 for left atrium, right ventricle, and right atrium.

### Tissue staining and microscopy

RNA *in situ* hybridization was performed using the RNAscope 2.5 High Definition and RNAscope Multiplex Fluorescent v2 assays from Advanced Cell Diagnostics, Inc. (catalog numbers 322370 and 323100, respectively) following the manufacturer’s protocols with the following modifications. During tissue preparation, fresh frozen tissue was sectioned at 15 um and mounted onto Superfrost Plus Slides (VWR). Fixation was performed in 4% PFA for 10 min at 4°C. Protease treatment was performed using RNAscope Protease III for 15 min at RT. Probes for *ADAMTS4*, *DCN* and *RYR2* were obtained from Advanced Cell Diagnostics. Hematoxylin and eosin or Masson’s Trichrome stains were performed on paraffin embedded sections according to standard protocols.

### Marker gene identification

Genes discriminating each cluster were identified by calculating the area under the receiver operating characteristic curve (AUC) for all genes comparing cells from the target cluster to all other cells not included in that cluster. The AUC will indicate how well a gene discriminates cells of a given cluster from those of all other clusters, with a value of 0.50 designating no discrimination and a values of 1 designating perfect discrimination. Two clusters determined to contain a high fraction of cytoplasmic material based on the proportion of exonic reads detected with scR-Invex were excluded from these calculations. Data were normalized by dividing the number of UMI for each transcript by the total UMI for the cell, multiplying by 10,000, and taking of the natural log of this result. The classifier used to calculate the AUC was built by taking the normalized expression values in each cell as predictions, and the cell cluster assignment as the class being predicted. Genes with an AUC greater than 0.70 and an average natural log fold-change > 0.6 were selected as markers of a cluster. These cutoffs were chosen to balance selecting genes with moderate discrimination for the cluster of interest, while not being overly inclusive so that a reasonably sized set of genes (hundreds for the larger clusters and a less than 10 for the smallest cluster) was available for each cluster. A less stringent AUC cutoff was required than others have used in the past [45, 46] as: 1) the reference group for a cluster of interest sometimes contains cells with similar expression profiles (e.g., multiple fibroblast clusters) and 2) single nuclei RNA-seq create a generally higher noise ratio than whole cell RNA-seq. In some clusters, these gene lists did not provide sufficient information to identify cell types. In those cases, genes expressed in > 5% of cluster cells that showed a standardized positive predictive value (PPV50) > 0.90 were also examined [47, 48]. For each gene, the PPV50 between the target cluster of interest and all cells from other clusters quantifies the probability that a cell is of the target cluster when it expresses that gene of interest (UMI > 0), standardizing to an equal number of cells between clusters (prevalence=50%). Standardization was necessary given the highly variable numbers of cells within each cluster, which systematically reduce the number of genes found with an unstandardized PPV for smaller clusters.

### Sub-clustering

To uncover potential sub-clusters of cells within the major clusters identified above, a simple sub-clustering procedure was performed for the follow cell types: cardiomyocytes (clusters 3, 4, 6, 15), fibroblasts (cluster 1, 2, 14), endothelial cells (cluster 9 and 10), pericytes (cluster 7), macrophages (cluster 8), adipocytes (cluster 11), vascular smooth muscle (cluster 13), neuronal (cluster 16), and lymphocytes (cluster 17). A new neighborhood graph was built for each of these groups using cosine distance based on the latent variables derived from the global scVI model. Louvain clustering was applied using varying resolution (ranging from 0.2 to 0.6) to establish new sub-groups. AUC was calculated for each sub-cluster compared to the remaining cells in the cluster. Genes with AUC greater than 0.65 and an average natural log fold-change > 0.5 were selected as markers of sub-clusters. A more liberal cutoff was employed here as cell sub-clusters look more similar to one another on average. Similarly to the global map, when necessary genes expressed in > 5% of target subcluster cells with a PPV50 > 0.90 were examined to help determine potential cell sub-types. Putative spurious sub-groups were identified based on an elevated proportion of exonic reads to all reads, which are often marked by increased mitochondrial genes.

### Differential gene expression analysis

Between-chamber and between-sex differential gene expression analyses were performed for the top five most abundant cell types in the aggregated four chamber map. This included cardiomyocytes (cluster 3, 4, 6, 15), fibroblasts (cluster 1, 2, 14), endothelial cells (cluster 9 and 10), pericytes (cluster 7), and macrophages (cluster 8). Additional sub-clusters within the cardiomyocytes and endothelial cells were removed if they had an enriched proportion of spliced transcripts, often accompanied by mitochondrial gene markers (see above).

Within each cell type, a generalized linear mixed model framework was employed using the R package lme4.[49] For a given gene in a given cell type, we first assumed that the UMI counts in cell *i* from experiment *j* of individual *k*, denoted *y_ijk_*, followed a negative binomial distribution,[50] 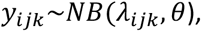 where θ represents inverse over-dispersion.[50] In many cases, θ approached infinity and we therefore reverted to a Poisson assumption, *y_ijk_* ∼*Poisson* (*λ_ijk_*),_’_, if θ > 10,000 for either the null or the full model. We constructed two generalized linear mixed models for log(*λ_ijk_*), specifically:

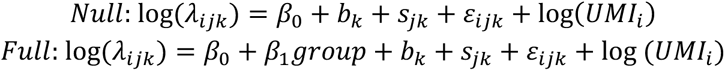

where *β*_o_ is a global mean UMI, *β*_1_ is the fixed effect for the group of comparison (chamber or sex), log(*UMI*_i_) is an offset of the total UMI in cell *i*, and *b_k_*, *s_jk_*, and ε*_ijk_* are random effects for biological sample, experiment and residual error normally distributed with mean 0 and variances 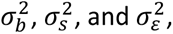 respectively. Any genes where θ < 0.10 from either the null or full negative binomial model were removed as very high over-dispersion created problems in model convergence.

In lme4 notation, the negative binomial mixed model was fit as:

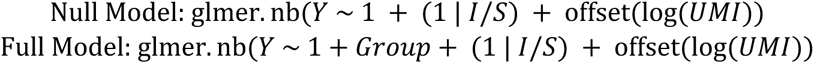

And the Poisson model was fit as:

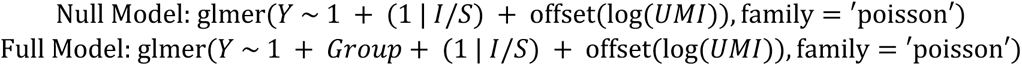

where *Y* represent UMI counts, *I* is a random effect of biological individual, *S* is a random effect of experiment, *UMI* are the total UMI counts in the given cell, and *Group* represents the fixed effect comparison of interest.

Significance was tested using a likelihood ratio test comparing the full model to the null model. Only genes expressed in at least 1% of either group in the given comparison were tested. To avoid capturing genes only present in the ambient background RNA or genes whose expression comes from cluster misclassification, only genes with a PPV50 > 0.55 or PPV50 > 0.50 for the cluster of interest were included for testing chamber comparisons and sex comparisons, respectively. To account for multiple testing in a given comparison of interest, a false discovery rate (FDR) correction using the Benjamini-Hochberg procedure was applied jointly for all genes tested across the five considered cell types. Any gene with an FDR corrected *P* < 0.01 was considered significant.

### Gene ontology analysis

Gene ontology analysis was performed using the R package GOstats version 2.46.0. Ensembl identifiers were mapped to Entrez gene identifiers when possible for compatibility with GOstats gene ontologies. The gene universe was set to all protein coding genes that were successfully mapped and only gene sets with a minimum size of 5 were considered for enrichment testing.[51] For each set of marker genes, a hypergeometric test was performed to test for enrichment of genes in each ontology, considering only ontologies with at least one gene overlapping the given marker list. A Bonferroni significance threshold was used for each set of markers correcting for the number of ontologies tested.

### Genome-wide association study integration

For six cardiometabolic traits with genome-wide association studies (GWAS), we looked for enriched heritability around marker genes of given cell types using stratified linkage disequilibrium (LD) score regression.[52, 53] We considered major cell types for this analysis, excluding low quality sub-clusters identified as described above. Only genes with a total of at least 10 counts across all cells were considered. Gene coordinates were used from the GRCh37 Ensembl reference to align with LD score regression methods. When genes from the GRCh38 Ensembl reference were not available in GRCh37 Ensembl reference, coordinates were lifted back using liftOver [54] when possible. In total, 25,968 genes were considered. For each cluster, a new set of marker genes were identified based on having at least some discrimination for the cluster of interest over other cell types (AUC > 0.55). Single nucleotide polymorphisms (SNPs) within 100 KB of any gene identified this way were annotated for LD score regression based on 1000G European individuals. The LD score regression model was run including the baseline annotations generated in *Finucane et al* 2015 [55] only considering high quality HapMap3 SNPs. The six GWAS traits used included atrial fibrillation,[26] PR Interval,[56] QT Interval,[57] coronary artery disease,[58] LDL,[59] and type 2 diabetes.[60] European ancestry-specific results were used when available to be most consistent with the LD reference panel.

Additionally, specific GWAS genes for atrial fibrillation, PR interval, QT interval, and coronary artery disease were highlighted based on colocalization of a genome-wide association signal and an expression quantitative trait loci (eQTL) from bulk sequence data of the relevant chamber of the heart. Colocalization was performed in a 1 MB region around the sentinel SNP of a GWAS locus using the coloc.abf function from the coloc package in R.[61] Allele frequency data was derived from the same European 1000 Genomes [62] samples used in the LD score regression analysis described above. Left ventricle eQTL data was taken from the Genotype-Tissue Expression (GTEx) project [25] based on 272 samples and left atrial eQTL data was taken from the MAGNet repository (http://www.med.upenn.edu/magnet/) based on 101 individuals.[26] Genes estimated to have a greater than 0.60 probability that the GWAS signal and eQTL signal share a causal variant we considered putative GWAS genes.

## DATA AVAILABILITY

Raw sequence data will be made available through dbGaP accession number phs001539.v1.p1. Processed data will be available through the Broad Institute’s Single Cell Portal under study ID SCP498.

## RESOURCES

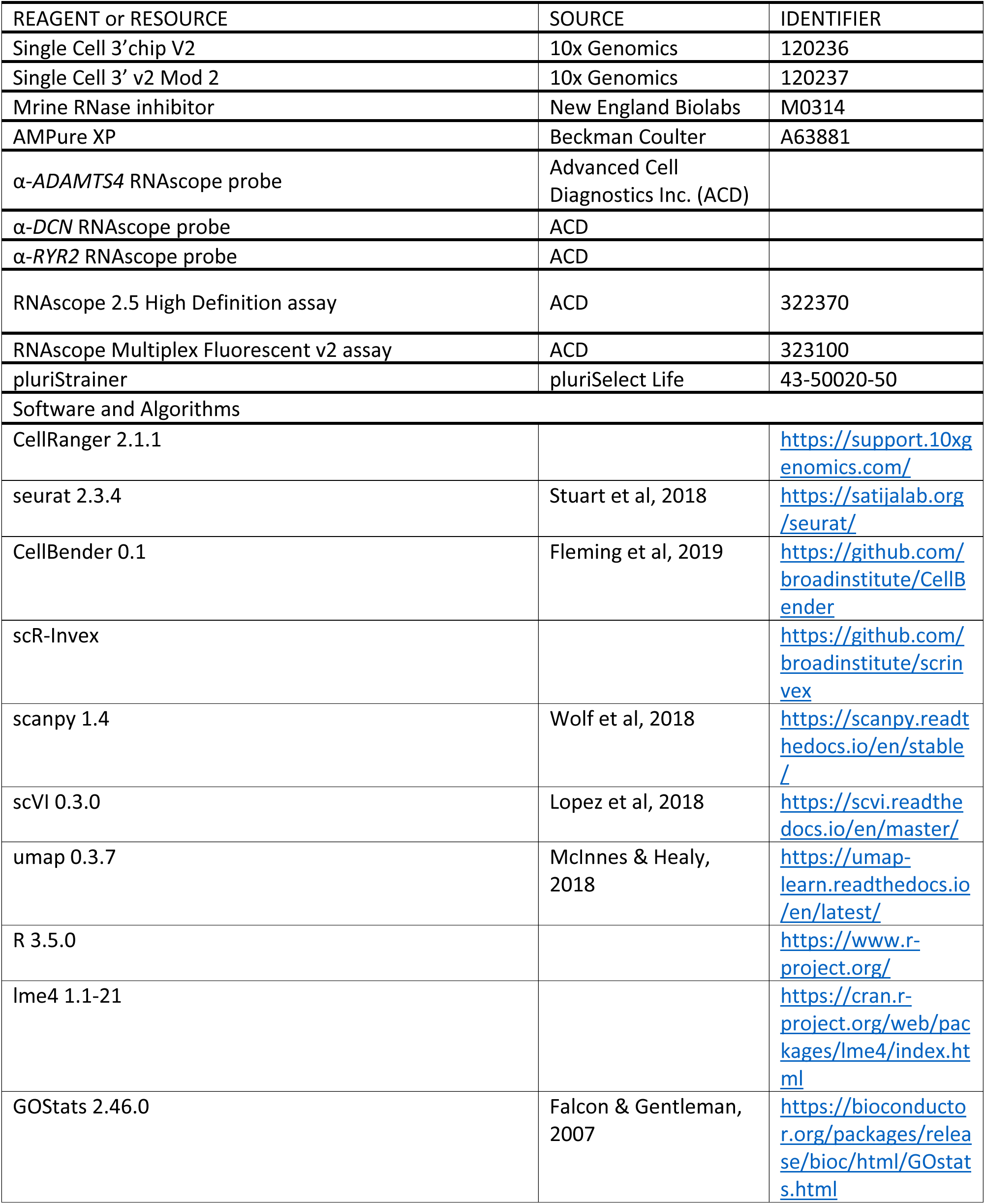
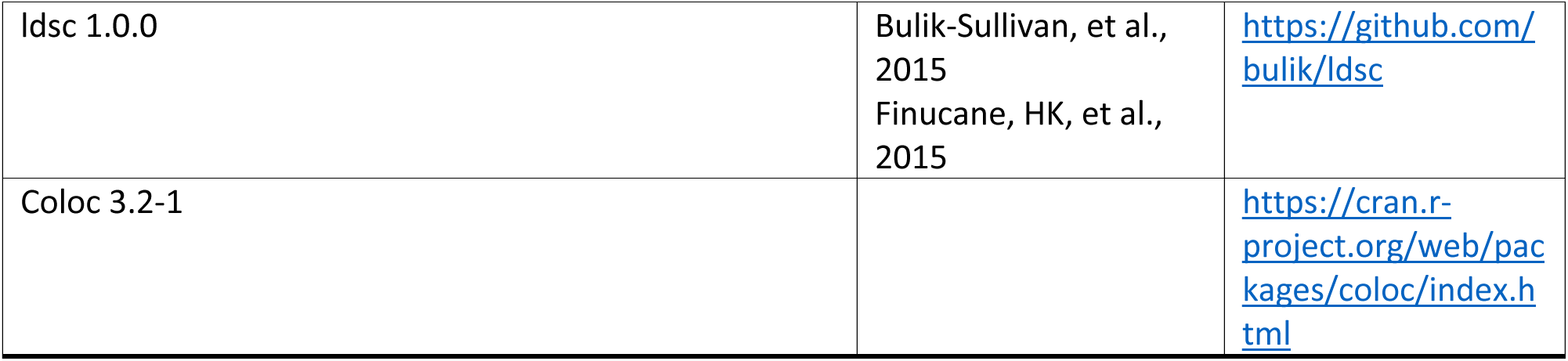

