## Supplemental Figures for "Transcriptional and Cellular Diversity of the Human Heart"

**Running Title:** Single cell transcriptomics of the human heart

**Word count:**

**Keywords:** Heart, single cell sequencing, cardiovascular disease, genetics

**Corresponding Author:**
Patrick T. Ellinor, MD, PhD
The Broad Institute of MIT and Harvard
75 Ames Street
Cambridge, MA 02142


**Table of Contents:**


| Description | Page |
|---|---|
| Supplemental Tables 1-5: provided as .xls files | 3 |
| Table ST1. Summary of quality control metrics for each sample processed. | |
| Table ST2. Marker genes for clusters identified in the joint UMAP plot. | |
| Table ST3. Marker genes for subclusters identified following secondary clustering of major cell types. | |
| Table ST4. Results of differential expression analysis for each chamber level comparison for cardiomyocytes, endothelium, fibroblasts, macrophages and pericytes. | |
| Table ST5. Results of differential expression analysis between males and females within all chambers, and each chamber individually, for cardiomyocytes, endothelium, fibroblasts, macrophages, and pericytes. | |
| Supplemental Figure S1: Analytic workflow and quality control metrics. | 4 |
| Supplemental Figure S2: Establishment of quality control metrics. | 5 |
| Supplemental Figure S3: Subclustering analysis of cardiomyocytes and pericytes. | 7 |
| Supplemental Figure S4: Histological analysis of donor tissue. | 8 |
| Supplemental Figure S5: Intersection of snRNAseq data with clinical testing panels and eQTL data. | 9 |

**Supplemental Tables**

**Tables ST1-ST5 are contained within the attached excel spreadsheets.**

**Figure S1. Analytic workflow and quality control metrics.** A: Analytic workflow for post-sequencing quality control through initial clustering. B: Table detailing the contribution of each sample to the final map by chamber and replicate number. UMI = unique molecular identifier; nGene = number of genes detected in a given cell; nUMI = the total UMI in a given cell; Q3 = 75$^{th}$ percentile; IQR = Interquartile Range; MT = Mitochondrial; RA = Right Atrium; LA = Left Atrium; RV = Right Ventricle; LV = Left Ventricle.

A

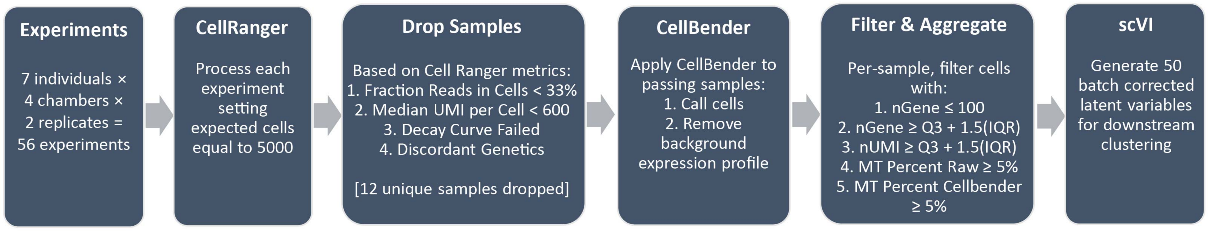


B

| RA (4 Female, 2 Male) | | | LA (3 Female, 3 Male) | | |
|---|---|---|---|---|---|
| RA_1221_1 | RA_1681_1 | RA_1708_1 | LA_1600_1 | LA_1681_1 | LA_1708_1 |
| | RA_1681_2 | RA_1708_2 | LA_1600_2 | LA_1681_2 | LA_1708_2 |
| RA_1600_1 | RA_1702_1 | RA_1723_1 | LA_1666_1 | LA_1702_1 | LA_1723_1 |
| RA_1600_2 | RA_1702_2 | RA_1723_2 | LA_1666_2 | LA_1702_2 | LA_1723_2 |
| **RV (3 Female, 2 Male)** | | | **LV (3 Female, 3 Male)** | | |
| RV_1221_1 | RV_1702_1 | RV_1723_1 | LV_1221_1 | LV_1681_1 | LV_1708_1 |
| RV_1221_2 | RV_1702_2 | RV_1723_2 | | LV_1681_2 | LV_1708_2 |
| RV_1666_1 | RV_1708_1 | | LV_1666_1 | LV_1702_1 | LV_1723_1 |
| RV_1666_2 | RV_1708_2 | | LV_1666_2 | LV_1702_2 | LV_1723_2 |

**Male:**
- P1702
- P1666
- P1681

**Female:**
- P1221
- P1600
- P1723
- P1708

**Figure S2. Establishment of quality control metrics.** A: UMAP plots generated both before and after batch correction by scVI. Colors of each dot correspond to the patient sample from which the cell arose. B: UMI decay curve from sample LV_1723_2 broken down by cell type after cell calling by CellBender. Colors correspond to the cell types as labeled. C: UMAP plot which displays the absolute percentage of mitochondrial (MT) reads in each cell. D: Density plot displaying the distribution of reads which map to exonic regions over total reads by cell type. Color corresponds to the cell clusters as called within the global Louvain clustering. Both clusters labeled as “cytoplasmic” indicate a higher than normal number of reads in the exonic regions.

**Figure S2:**

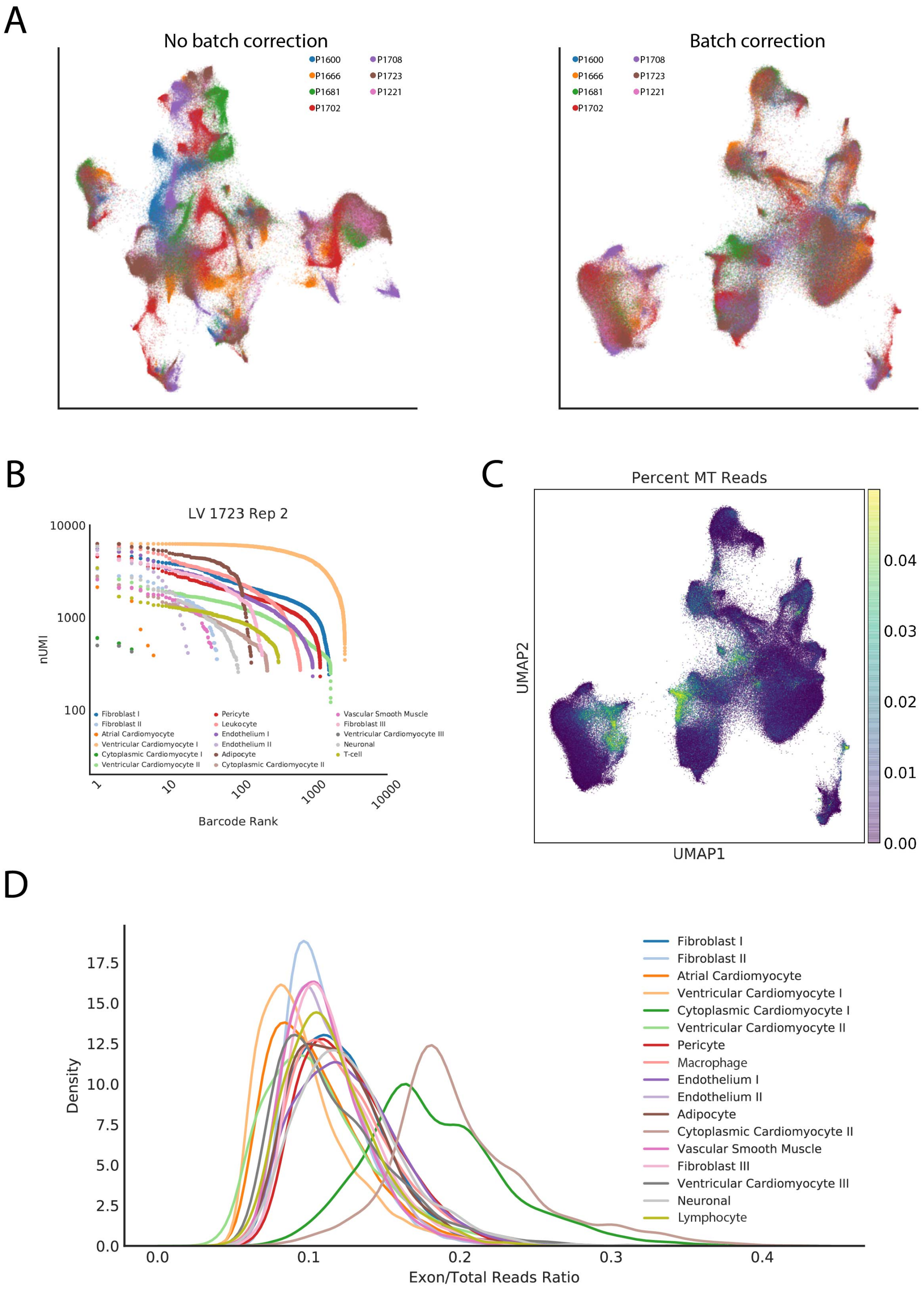
A
No batch correction
P1600
P1666
P1681
P1702
P1708
P1723
P1221
Batch correction
P1600
P1666
P1681
P1702
P1708
P1723
P1221
B
LV 1723 Rep 2
10000
1000
100
nUMI
Fibroblast I
Fibroblast II
Atrial Cardiomyocyte
Ventricular Cardiomyocyte I
Cytoplasmic Cardiomyocyte I
Ventricular Cardiomyocyte II
Pericyte
Leukocyte
Endothelium I
Endothelium II
Adipocyte
Cytoplasmic Cardiomyocyte II
Vascular Smooth Muscle
Fibroblast III
Ventricular Cardiomyocyte III
Neuronal
T-cell
1
10
100
1000
10000
Barcode Rank
C
Percent MT Reads
UMAP2
UMAP1
0.04
0.03
0.02
0.01
0.00
D
17.5
15.0
12.5
10.0
7.5
5.0
2.5
0.0
Density
0.0
0.1
0.2
0.3
0.4
Exon/Total Reads Ratio
Fibroblast I
Fibroblast II
Atrial Cardiomyocyte
Ventricular Cardiomyocyte I
Cytoplasmic Cardiomyocyte I
Ventricular Cardiomyocyte II
Pericyte
Macrophage
Endothelium I
Endothelium II
Adipocyte
Cytoplasmic Cardiomyocyte II
Vascular Smooth Muscle
Fibroblast III
Ventricular Cardiomyocyte III
Neuronal
Lymphocyte

**Figure S3. Subclustering analysis of cardiomyocytes and pericytes.** Results of major cell type subclustering for cardiomyocytes (A) and pericytes (B). Left panel displays the distribution of the identified subclusters within the global UMAP plot for all chambers. Each dot represents a cell, colored by its respective subcluster. Center panel is a density plot which displays the ratio of reads which lie in exonic regions as a percentage of total reads in each subcluster. Right panel is a dot plot displaying the genes with the highest AUC for each subcluster or those which are specifically mentioned within the text (*KCP*). The size of the dot represents the percentage of cells in the cluster in which each gene is detected and the color reflects the mean $\log_2$ expression. CM: Cardiomyocytes, PC: Pericytes.

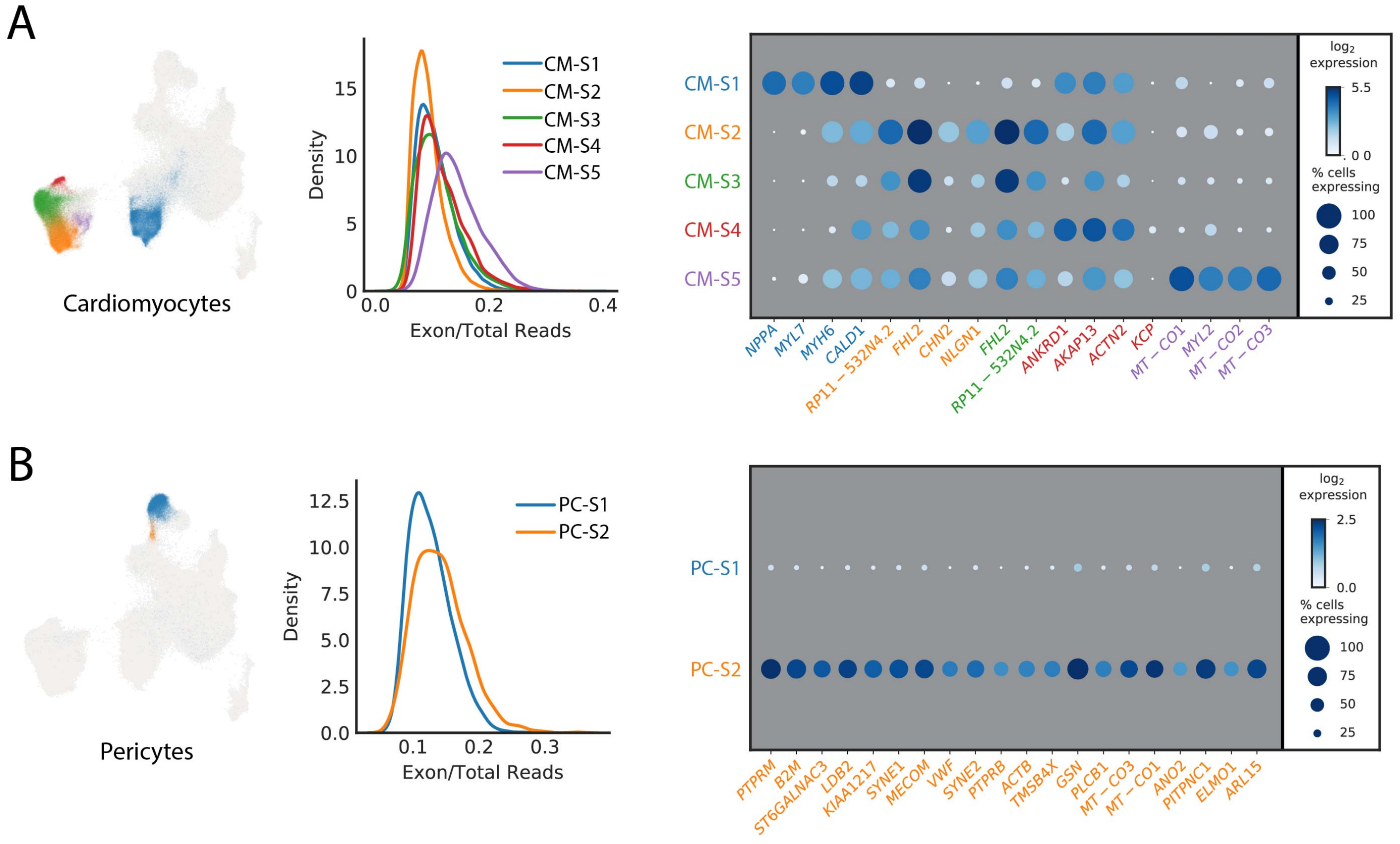

**Figure S4. Histological analysis of donor tissue.** A: Masson's Trichrome staining of representative regions from all four chambers in patient P1600. Scale bar indicates 1mm. B: Hemotoxylin/eosin staining of samples from patient P1723. Scale bar represents 1mm. Arrows in right panel highlight regions of myocardial adiposity. RA: Right Atrium, LA: Left atrium, RV: Right ventricle, LV: Left ventricle.

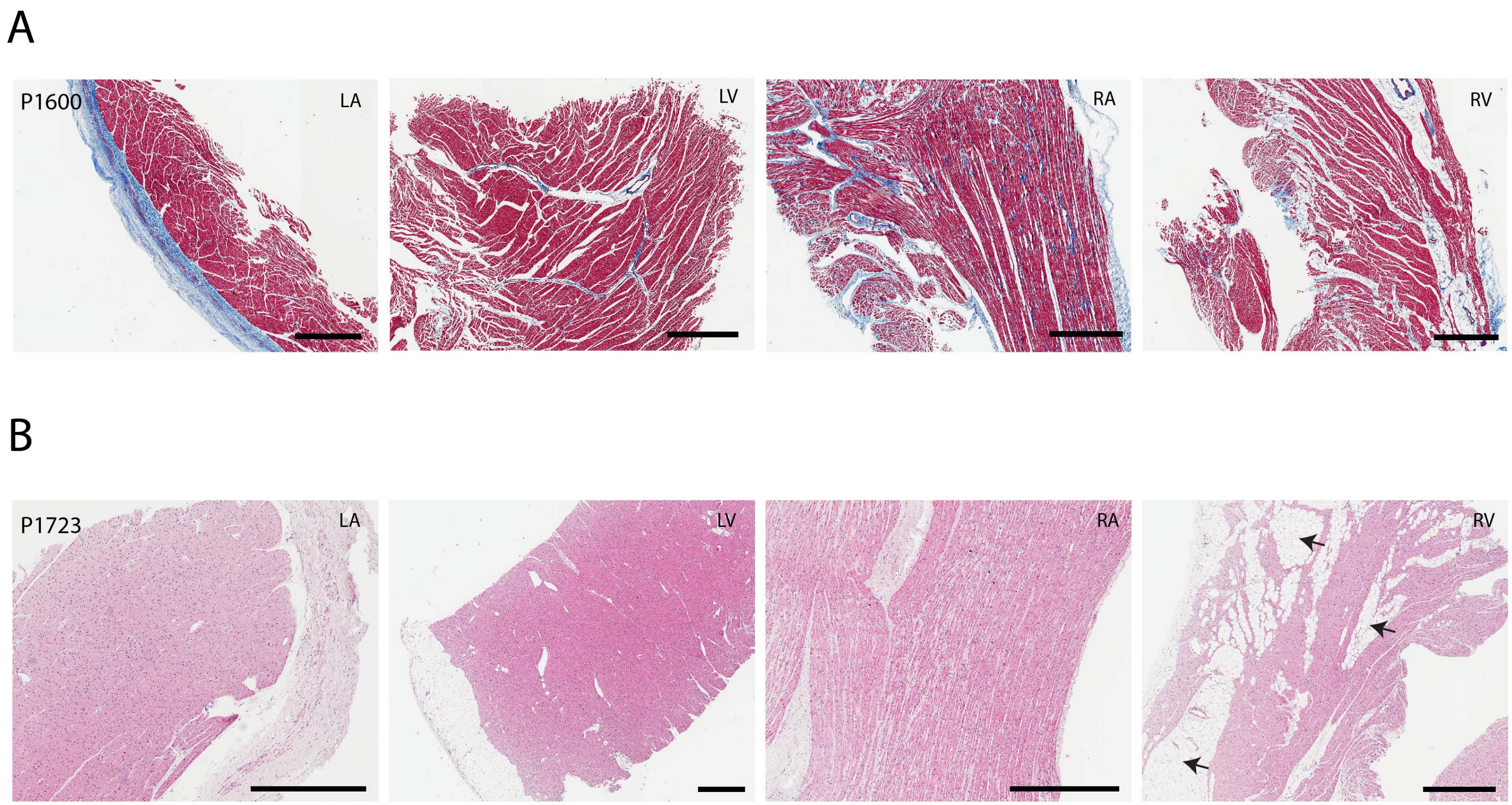

**Figure S5. Intersection of snRNAseq data with clinical testing panels and eQTL data.** A: Dot plot for genes currently on standard arrhythmia clinical testing panels. The size of each dot represents the percent of cells in which the gene of interest is detected and the shading represents the relative expression of the gene. Color of the genes correspond to the cell type for which the AUC reaches 0.70 or greater. Genes with black color indicate no cell type which reaches this threshold. Size and shade of the dot corresponds percentage of cells and relative expression, respectively. B: Dot plots for genes identified for eQTL analysis of left atrial and left ventricular tissue. SNPs used for eQTL analysis are those derived from the genome-wide association studies listed for cardiovascular traits and diseases.

**Figure S5:**

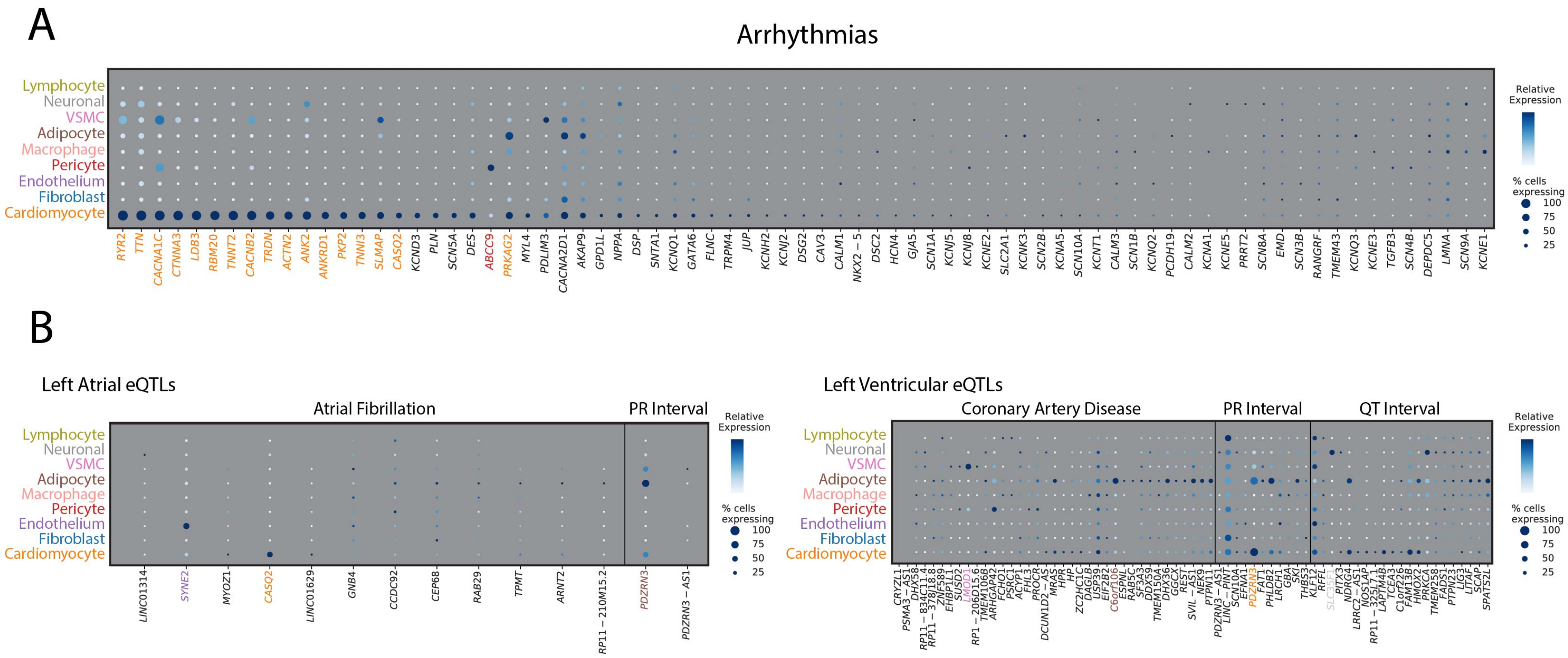
A
Arrhythmias
Lymphocyte
Neuronal
VSMC
Adipocyte
Macrophage
Pericyte
Endothelium
Fibroblast
Cardiomyocyte
Relative Expression
% cells expressing
100
75
50
25
B
Left Atrial eQTLs
Atrial Fibrillation
PR Interval
LINC01314
SYNE2
MYOZ1
CASQ2
LINC01629
GNB4
CCDC92
CEP68
RAB29
TPMT
ARNT2
RP11 – 210M15.2
PDZRN3
PDZRN3 – AS1
Left Ventricular eQTLs
Coronary Artery Disease
PR Interval
QT Interval
